# A multiomic lifespan signature in genetically diverse, diet-restricted mice

**DOI:** 10.64898/2026.05.08.723574

**Authors:** Johanna Y. Fleischman, Celeste Sandoval, Ngoc Vu, Martin Mullis, Phillip Seitzer, Leanne J.G. Chan, Niclas Olsson, Thao Nguyen, Aleksandr Gaun, Alison Luciano, Jonathan O’Brien, Jenny Vu, Laura Robinson, Andrea Di Francesco, Wenzhou Li, Sean R. Hackett, Rob Keyser, Fiona E. McAllister, Gary A. Churchill, Bryson D. Bennett

## Abstract

Dietary restriction extends lifespan across model organisms, but the plasma molecular changes mediating this effect remain incompletely characterized. We present a longitudinal multiomic analysis of 2,234 plasma samples from 960 Diversity Outbred mice subjected to intermittent fasting or caloric restriction and followed to natural death. Using mass spectrometry, we quantified 1,512 metabolites, lipids, and proteins and mapped their associations with diet, age and longevity. DR-induced molecular changes scale with caloric intake and modulate inflammatory, lipid catabolism, and oxidative stress pathways. Aging showed a biphasic signature with sharp acceleration beyond 85% of lifespan, demarcating terminal decline. Mediation and survival modeling both identified superoxide dismutase (SODE) and vascular cell adhesion molecule (VCAM1) as top lifespan predictors. Genetic analysis revealed 9,599 QTL, nine of which coincided with previously identified lifespan QTLs, and were largely related to immune regulation. These findings provide a rich multiomic and genetic resource for the aging research community.

## Main

Aging is a complex process and a major risk factor for many diseases. Dietary restriction (DR), a reduction in food intake without malnutrition, is one of the most robust non-genetic interventions shown to extend lifespan and improve healthspan across many model organisms^1^. DR delays all aging hallmarks including reduced nutrient signaling cascades, increased autophagy, and decreased inflammation^1,2^, but there is a lack of data on the effect of long-term DR in humans without malnutrition^3^. However, aging hallmarks are highly conserved across species^4^ and DR consistently improves primate metabolic and immune health^5–7^.

Dietary interventions that benefit health and longevity in rodents vary and include time-restricted feeding (TRF), intermittent fasting (IF), and chronic caloric restriction (CR)^2,8,9^. The Dietary Restriction in Diversity Outbred mice (DRiDO) study was a 5-year study of 960 female genetically diverse Diversity Outbred (DO) mice subjected to two CR and two IF regimens and followed to natural death^10–12^. Previous analyses of lifespan, body weight, and blood cell distributions from the DRiDO study found that lifespan and reduced adiposity are correlated with caloric intake, and that improved health and extended lifespan are not synonymous^12,13^. Within DR groups and independent of diet, resilience to weight loss during periods of stress was associated with longer lifespan. Here, we present a multiomic analysis of plasma samples from the DRiDO study and highlight metabolites, lipids, and proteins affected by DR, genetics, and aging. In mice and humans, DR has previously been associated with changes in lipid and amino acid metabolites^14,15^ and apolipoproteins^16,17^, but it remains unclear how metabolites, lipids, and proteins collectively change over the full course of aging or in response to DR. No studies have directly compared changes in the plasma multiome between IF and CR over the lifespan, nor examined whether different DR modalities converge on shared molecular mechanisms of longevity.

This paper presents the largest multiomic mass spectrometry molecular profiling study in naturally-aged genetically heterogeneous mice, as most previous studies have examined either single molecular classes or cross sectional timepoints. We perform an unbiased multiomic analysis of longitudinal plasma samples of DO mice and reveal a robust biphasic aging signature that parses healthy aging from end-life. Normalizing trajectories to the proportion of life lived is a particularly powerful approach, as it accounts for the substantial variation in lifespan across individuals and dietary groups, and enables the identification of molecular changes that track with biological rather than chronological age. This analysis reveals distinct aging phases and conserved plasma signatures that predict individual lifespan and illuminate candidate mediators linking DR to longevity. Further, we were powered to identify quantitative trait loci (QTL) associated with immune-related lifespan phenotypes and provide an in-depth resource of thousands of QTLs of molecular concentrations in blood plasma.

## Results

### Lifespan-monitored multiomic study yields 1512 traits for diet, aging, and genetic analysis

This study simultaneously captures genetic diversity, lifespan, healthspan, and longitudinal multiomic molecular profiling, and provides a comprehensive resource for investigating the plasma molecular landscape of aging and DR. A total of 960 Diversity Outbred female mice were assigned to one of five dietary regimens at 6 months of age: ad libitum (AL), intermittent fasting (IF) 1 or 2 days per week (IF-1D and IF-2D), or caloric restriction (CR) designed to restrict intake by 20% or 40% less than the AL group (CR-20 and CR-40) (Fig. 1a). Plasma samples were collected longitudinally at 6 months (Year 1), 16 months (Year 2), and 28 months (Year 3) of age from mice that survived to each respective timepoint; mice that died prior to the first blood draw were excluded from all analyses (Fig. 1b). Animals lived out their natural lifespan, and mean lifespan increased proportionally with reduction in caloric intake, with the actual decrease in calories calculated to be +3%, -12%, -27%, and -44% for the IF-1D, IF-2D, CR-20, and CR-40 groups compared to the AL group, as described previously^12^ (Fig. 1c). Mice were enrolled in 12 waves from March 2016 to November 2017 to minimize the confounding of seasonal changes^12^.

**Fig. 1:**
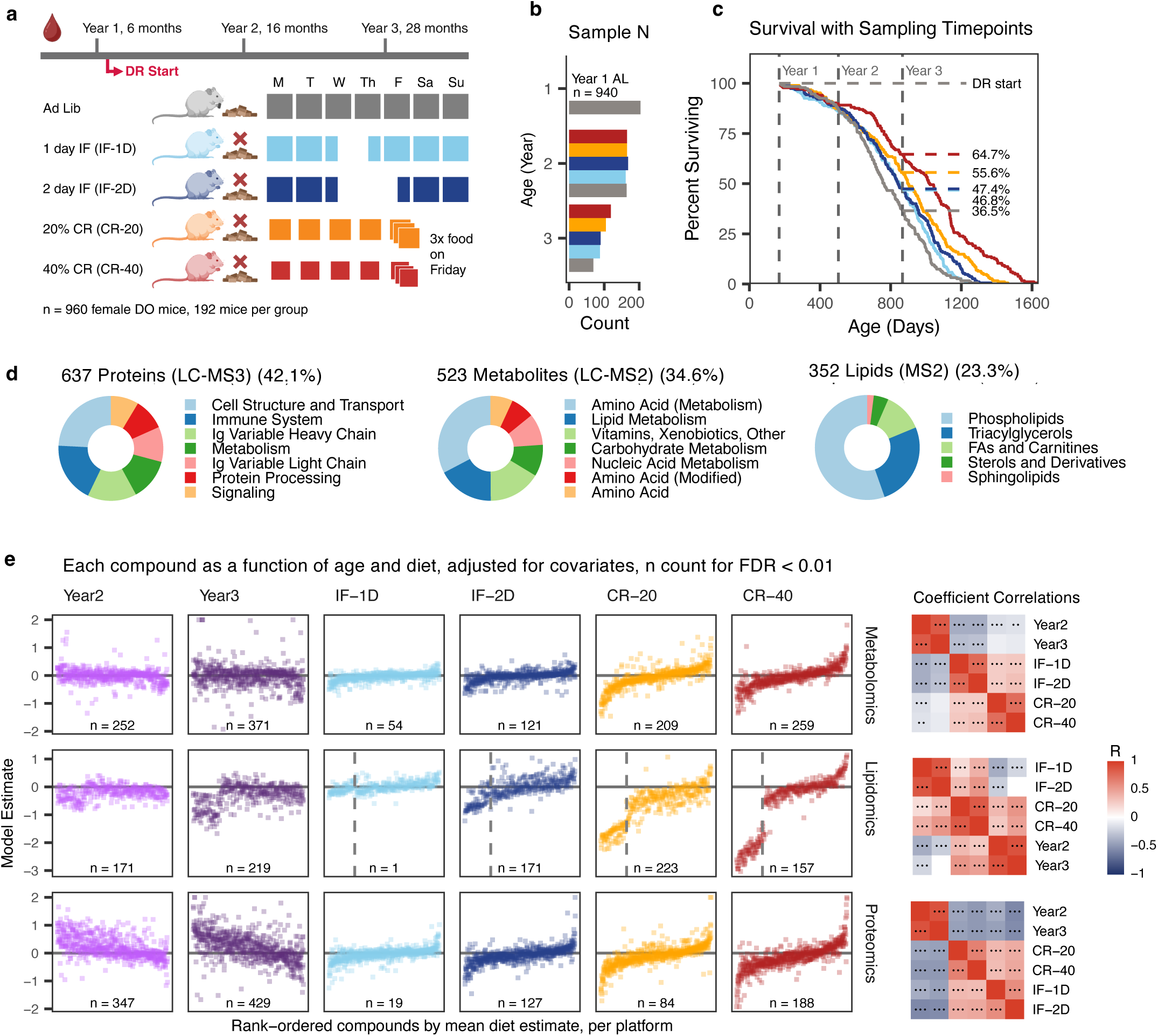
Lifespan monitored multiomic study yields 1512 traits for diet, aging, and genetic analysis. **a,** Female Diversity Outbred mice underwent one of five dietary restriction (DR) regimens at 6 months of age, and plasma samples were collected before intervention (Year 1, n = 940), one year after intervention start (Year 2, n = 823), or two years after intervention start (Year 3, n = 471). **b,** Sample distribution by diet and timepoint. **c,** Mice under DR lived longer than AL counterparts. **d,** Mass spectrometry data acquisition resulted in 1,512 annotated traits across all data types, with traits manually assigned into broad categorical classifications (Ig = immunoglobulin, Supp. Table 1). **e,** Diet-associated compounds were determined by quantifying main effect diet terms, adjusted for age, bodyweight, and sample collection weekday. Compounds for shown model terms are arranged by mean diet term estimate for each data type. The number of significant compounds is determined based on an FDR < 0.01 threshold. Dotted line within lipidomics indicates that compounds left of line are all triglycerides. Correlates between Diet and Age estimates are shown on the right. •••/••/• = pvalue < 0.001, 0.01, 0.05. Model estimates are floored at -2 and 2.

**Table 1.** Pre and Post Extraction standards for Metabolites and Lipids.

| method | pre/post-extraction | Chemical name | Vendor | concentration (ug/mL) |
| --- | --- | --- | --- | --- |
| Metabolomics | pre | 2-Oxoglutarate-13C5 | Sigma Aldrich | 2 |
| Metabolomics | pre | Creatinine-d3 | Cayman Chemical | 2 |
| Metabolomics | pre | Hypoxanthine-13C5 | Cambridge Isotope Laboratories | 2 |
| Metabolomics | pre | L-Glutamine-c13 | Cambridge Isotope Laboratories | 2 |
| Metabolomics | pre | Malate-13C | Sigma Aldrich | 1.6 |
| Metabolomics | pre | Ornithine-d6 | Sigma Aldrich | 5 |
| Metabolomics | pre | Pyruvic acid-13C3 | Sigma Aldrich | 8 |
| Metabolomics | pre | Trimethylamine N-oxide-d9 | Cayman Chemical | 0.5 |
| Metabolomics | pre | algal labeled AA (500 mg) | Sigma Aldrich | 3.4 |
| Metabolomics | pre | Citrate-d4 | Sigma Aldrich | 4 |
| Metabolomics | pre | D-Glucose-13C6 | Sigma Aldrich | 4 |
| Metabolomics | pre | Fumaric acid-d4 | Sigma Aldrich | 5 |
| Metabolomics | pre | L-Asparagine-15N2 | Cambridge Isotope Laboratories | 5 |
| Metabolomics | pre | L-Carnitine-d3 | Sigma Aldrich | 2 |
| Metabolomics | pre | Lactic acid-13C3 | Sigma Aldrich | 4 |
| Metabolomics | pre | Succinic acid-13C4 | Sigma Aldrich | 4 |
| Metabolomics | pre | URACIL-15N2 | Sigma Aldrich | 1 |
| Metabolomics | post | 1,4-diaminobutane-D4 | Sigma Aldrich | 1 |
| Metabolomics | post | BENZOIC ACID-d5 | Sigma Aldrich | 0.1 |
| Metabolomics | post | Caffeine-13C3 | Cayman Chemical | 1 |
| Metabolomics | post | HEXANOIC ACID-d3 | Cayman Chemical | 0.1 |
| Metabolomics | post | L-Tryptophan-13C15N | Sigma Aldrich | 1.5 |
| Lipidomics | post | FA(C16)-d13 | Cayman Chemical | 0.31 |
| Lipidomics | post | PA(15:0-d7/18:1) | Avanti Research | 0.31 |
| Lipidomics | post | HexCer(d18:1-d7/15:0) | Avanti Research | 0.31 |
| Lipidomics | post | PE(p-18:0/18:1-d9) | Avanti Research | 0.31 |
| Lipidomics | post | PC(p-18:0/18:1-d9) | Avanti Research | 0.31 |
| Lipidomics | post | Carn(24:0)-d4) | Avanti Research | 0.31 |
| Lipidomics | post | EQUISPLASH mix | Avanti Research | 0.31 |
| Lipidomics | post | Cholesterol-d7 | Avanti Research | 7.8 |

Using mass spectrometry-based LC-MS/MS methodologies, we determined the relative concentrations of 523 metabolites across two methods (99 found in both ionization modes), 352 lipids, and 637 proteins from 2,234 plasma samples (Fig. 1d), a molecular scope substantially broader than prior longitudinal multiomic aging studies⁶⁴. Identified metabolites spanned a wide range of pathways, including amino acid, nucleic acid, and carbohydrate metabolism. The detected lipids covered 13 classes, with the majority of detected features being triacylglycerols (TGs) or phosphatidylcholines (Fig. 1d, Supp. Table 1). Tandem mass tag based proteomics identified and measured proteins across many classes, including proteins located in lipoprotein particles, immune-related signaling molecules, and both heavy and light chain variable immunoglobulin regions, which comprised 26% of the annotated protein traits in this analysis (Fig. 1d, Supp. Table 1).

### DR drives dose-dependent and convergent molecular changes that partially reverse the plasma signature of aging

We first examined the molecular changes induced by aging (Age) and DR (Diet), adjusting for bodyweight at collection timepoint, sample collection timing, and repeated measures from the same mouse (Fig. 1a, Ext. Fig. 1). Shifts in compound abundance scaled with caloric intake, as only 5% of compounds significantly (FDR < 0.01) differed from the AL abundances in the IF-1D group compared to 40% in the CR-40 group (Fig. 1e, Supp. Table 2). Triglycerides (TGs) were severely depleted in both CR groups, which may reflect either reduced dietary fat intake or increased fatty acid metabolism (Fig. 1e, left of dotted line). Among the 503 compounds significant in at least two DR interventions, 86% changed in the same direction regardless of restriction through IF or CR, suggesting that IF and CR converge on a shared molecular program rather than activating distinct biological pathways. Age-related molecular shifts intensified over time, with 51% of features significantly changed at Year 2 and 68% at Year 3 (Fig. 1e).

We separately examined the compounds that changed most significantly for Diet and Age terms using a Fisher combined test. Amongst the top 70 traits for each of Diet and Age, twelve overlapped and predominantly showed anticorrelation of effect, as compounds that increased with age tended to be depleted by DR, and compounds that declined with age tended to be enriched by DR (Fig. 2a). This directional opposition, observed across both innate immunity and TCA cycle metabolism, may represent a partial reversal of the biological aging signature by DR.

**Fig. 2:**
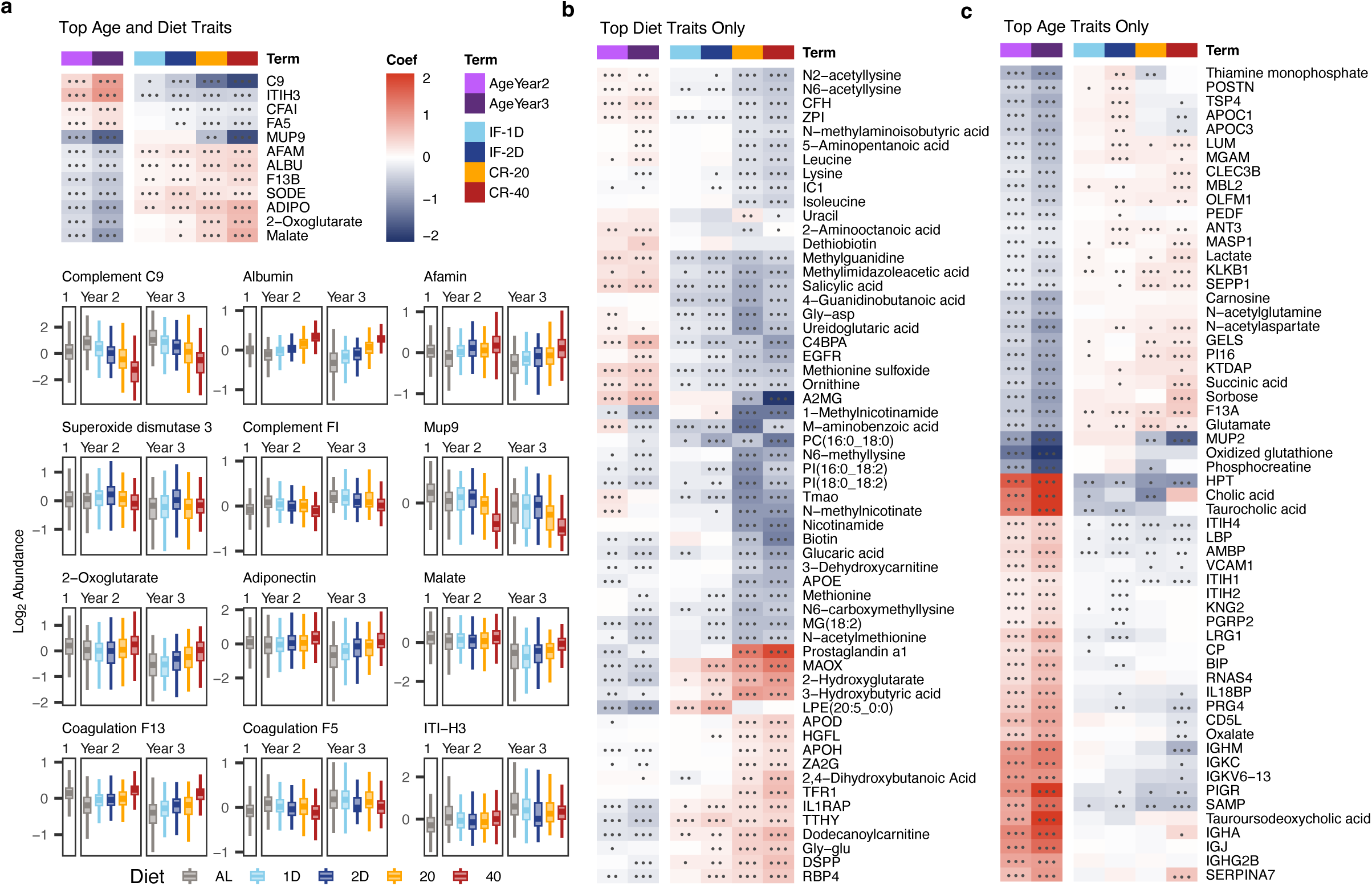
Circulating compounds are consistent between CR and IF and scale with age and caloric intake. **a,** Amongst the top 70 ranked terms for *Age* and *Diet* by lowest Fisher combined p-value on FDRs, 12 compounds were top ranked for both terms. Model estimates are shown in heatmap. Normalized sample values are shown in box plots. **b,** Heatmap of the remaining top-70 *Diet-*associated compound model estimates. **c,** Heatmap of the remaining top-70 *Age-*associated compound model estimates. •••/••/• = FDR < 0.001, 0.01, 0.05 on effect estimates. Model estimates are floored at -2 and 2.

Of the remaining 70 top-ranked compounds for Diet but not Age, many were amino acids or modified amino acids that were lowered by DR, which possibly reflects decreased nutritional intake (Fig. 2b). DR resulted in decreased levels of the chylomicron component APOE, consistent with the observed decrease in TGs (Fig. 1e). DR also increased APOH and APOD, which may reflect decreased inflammation and increased fatty acid beta-oxidation (Fig. 2b). We found lower levels of many members of the complement system: CO9, CFAI, CFAH, IC1, and C4BPA (Fig. 2a-b), which points to a broad downregulation of the complement cascade rather than modulation of the activity of individual components. This is consistent with recent studies in humans showing that CR reduces inflammaging via complement deactivation^18^. DR robustly elevated adiponectin (ADIPO) across all four interventions, positioning it as a candidate hormonal mediator of DR-associated longevity in this cohort. Adiponectin, which declined with age (Fig. 2a), is known to improve insulin sensitivity, reduce inflammation, and regulate energy metabolism by regulating lipid clearance and activating pro-metabolic pathways AMPK and PPARα^19–21^. DR also led to increases in the TCA cycle intermediates 2-oxoglutarate and malate (Fig. 2b), consistent with a phenotype of enhanced oxidative metabolism under CR.

The top traits associated with Age included immunoglobulin components (IGHG2B, IGHA, IGKC, IGHM) and proteins known to bind to Ig complexes (IGJ, PIGR) as well as bile acids (taurocholic acid, tauroursodeoxycholic acid, cholic acid) (Fig. 2c). The progressive increase in multiple bile acid species with age points to remodeling of hepatic metabolism and gut microbiome composition, which are increasingly being recognized as central to aging biology^22^. Compounds that decreased over time included lactate, succinate, and apolipoproteins (APOC1, APOC3), consistent with a broad decline in core metabolic activity with age.

### The plasma multiome captures a dual signature of healthy aging and terminal decline

The Age term in the preceding analysis captures molecular differences between fixed timepoints but cannot account for the substantial variation in lifespan across individuals and dietary groups. To resolve the aging trajectory while accounting for differential lifespans, we converted age to the proportion of life lived (PLL), using Diet as a covariate. We did the same with previously reported data from the DRiDO study, collected at separate timepoints^12^, including flow-cytometry quantified immune, metabolic cage, and body composition traits. We excluded pre-intervention samples that may not reflect the molecular aging profile of mice subsequently placed on DR.

PLL fits determined from a generalized additive model found 887 compounds with significant non-zero slopes (FDR < 0.05, Fig. 3a, Supp. Table 3). Similar trajectories were clustered using k-means into 10 groups as determined from a gap statistic (Fig. 3a, Ext. Fig. 2a). Most clusters showed a clear slope change in the last quarter of PLL, confirming two distinct phases of aging across diverse molecular classes (Fig. 3a, Ext. Fig. 2b). We determined the point of maximum curvature (PMC, the point with the greatest rate of slope change) over PLL, and determined the median PMC to be at 0.85 PLL, corresponding to an average of 116 days in the AL group up to 147 days in the CR-40 group (Fig. 3b).

**Fig. 3:**
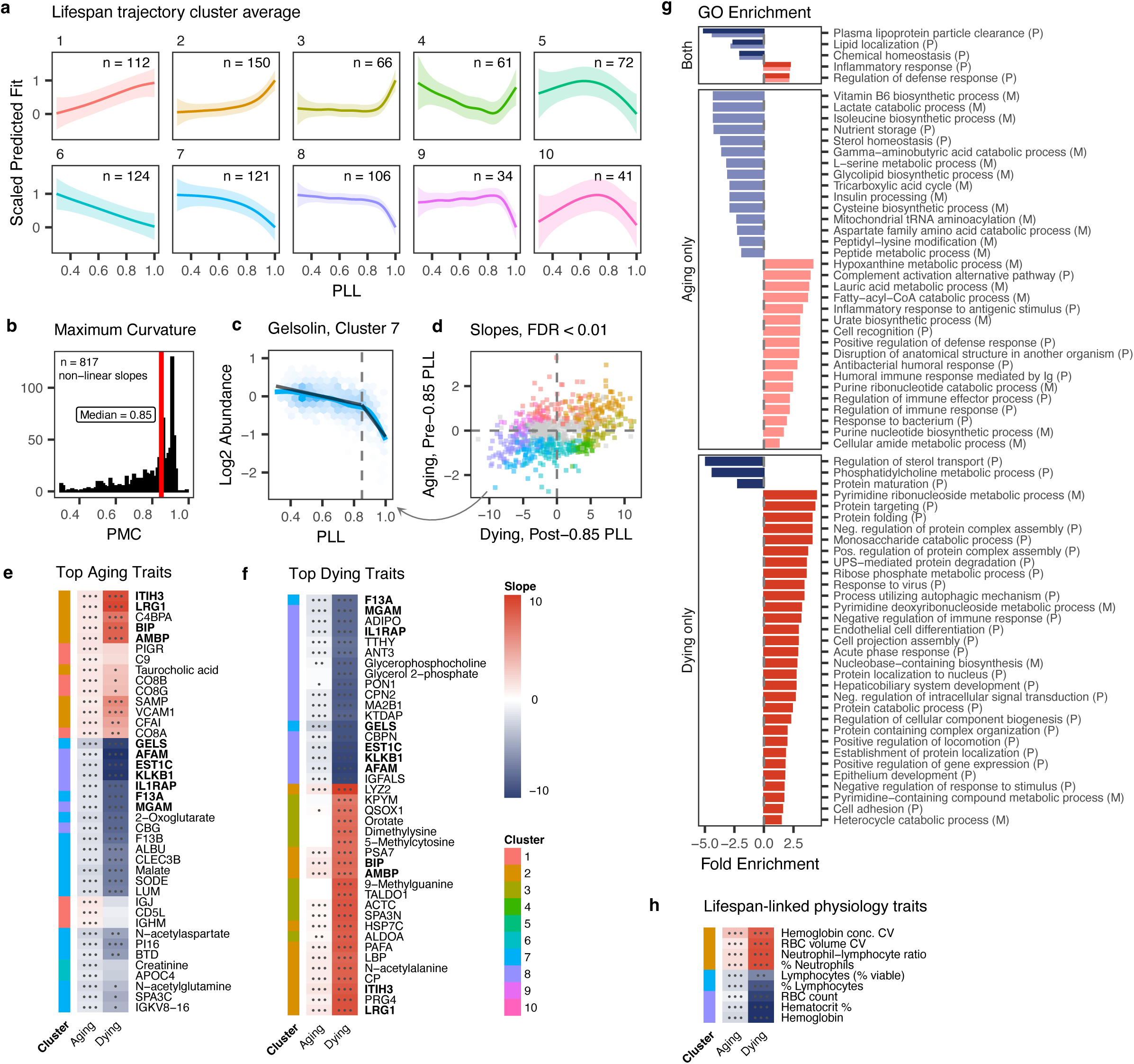
The plasma multiome captures a dual signature of healthy aging and terminal decline. **a,** Average non-linear aging trajectories as a function of Proportion of Life Lived (PLL) show distinct biphasic aging signature. **b,** The median point of maximum curvature (PMC) as the absolute maximum value of the second derivative of the GAM slope was 0.85 PLL. **c,** Colored lines show the GAM non-linear fit for PLL, and black lines represent slope coefficients for Aging and Dying as a function of PLL for Gelsolin (lowest Fisher combined FDR on Aging and *Dying* slopes). **d,** 593 traits had a significant non-linear slope for either Aging or Dying, calculated from linear fits on pre- and post-0.85 PLL data. 193 traits had both a significant Aging and Dying slope. Traits with lowest FDR (n = 40) using slopes from z-scored trait data are shown for Aging (**e**) and Dying (**f**) where slopes in both sets are bolded. **g,** Proteins and metabolites with significant (FDR < 0.01) coefficients for Aging or Dying were separated by enrichment or depletion. Pathways for GO biological process are shown when pathway FDR < 0.1, with redundant pathways removed. **h,** Aging and Dying slopes on z-scored data from blood traits mostly highly associated with lifespan. •••/••/• = FDR < 0.001, 0.01, 0.05 on effect estimates.

We next calculated separate linear slopes from pre- and post-0.85 PLL data points, termed Aging and Dying respectively (Ext. Fig. 2a). Linear and non-linear slopes were consistent, as exemplified by Gelsolin (lowest Fisher combined FDR for both slopes, Fig. 3c). We found 471 significant slopes for Aging and 320 for Dying, of which 198 were significant for both (Fig. 3d). Of those significant for both Aging and Dying, 81% were directionally consistent, with Dying slopes typically much steeper than Aging slopes, suggesting that during terminal decline there is an acceleration of at least some of the changes existent during healthy aging.

Of the 40 Aging and Dying traits with the lowest FDR (Fig. 3e-f), eleven were shared between the two phases and were related to the acute-phase inflammatory response (bolded traits, Supp. Table 4). To better understand the phase-specific pathways, we performed GO enrichment on significant proteins (“P”) and metabolites (“M”) (Fig. 3g, Supp. Table 5). Shared Aging and Dying pathways included lipoprotein particle depletion and enrichment of inflammatory response proteins. Strikingly, complement and humoral activation were unique to the Aging phase (Fig. 3g, Fig. 3e, IGJ, PIGR, CO9), indicating that innate and adaptive immune remodeling is a hallmark of healthy aging that may be extinguished by the onset of terminal decline. Aging was further marked by enrichment of purine metabolites and depletion of amino acid and TCA cycle components (Fig. 3g), consistent with a progressive dysfunction in metabolism during aging beyond the regulation of lipid homeostasis. Similar to many of the molecular measures, we also observed a biphasic pattern of slower Aging followed by much steeper Dying slopes in the hemoglobin, neutrophil, and lymphocyte levels previously identified as the blood cell traits most strongly correlated with lifespan in this cohort^12^ (Fig. 3h).

In addition to an apparent acceleration of the aging process during Dying, we also found a distinct set of pathways that were non-overlapping with Aging pathways, including the enrichment of protein and nucleobase catabolic processes (Fig. 3g). This pattern is consistent with active cellular disintegration and the release of intracellular contents into the circulation, a process that does not appear to occur in healthy aging. Together, these results suggest that Dying is both an acceleration of the Aging process and reflects cellular breakdown specific to end-life.

### Lifespan extension is associated with both DR-dependent and independent compounds

To determine what molecular traits best predicted the longevity of an individual mouse, we analyzed Year 2 samples, excluding those from the final 0.15 of PLL, to omit samples from the dying phase. We employed two complementary analytical approaches: first, mediation analysis to identify which traits frequently integrated the pairwise signal between other compounds and lifespan as molecular hubs; and second, a Lasso-penalized Cox survival model to identify which traits best predicted remaining lifespan while accounting for covariation. The convergence of both approaches on the same proteins provides the strongest evidence for their biological relevance.

The union of top compounds from both approaches showed a variety of patterns: many were affected by both Diet and Aging, but some were not. Of those affected by at least one Diet and Aging, 75% had opposite directionality between Diet and Aging, signaling potential reversal of effects (Fig. 4a). In the mediation analysis, most of the significant interactions involved one of a few compounds, including leucine-rich glycoprotein (LRG1, n = 56), superoxide dismutase 3 (SODE, n = 52), albumin (ALBU, n = 50), and vascular cell adhesion molecule 1 (VCAM1, n = 35) (Fig. 4b, Supp. Table 6), identifying these proteins as hubs through which the largest number of other molecular traits relate to lifespan. SODE and ALBU are both significant across all DR interventions as top Age and Diet traits (Fig. 2a) whereas LRG1 and VCAM1 are less associated with DR but highly associated with Aging (Fig. 3e), suggesting that these proteins show both DR-dependent and independent longevity signal. Direct DR-lifespan mediation was underpowered, but compounds significant at nominal p-values reflect those affected by both Aging and DR (Ext. Fig. 3, Fig. 2a, Supp. Table 7).

**Fig. 4:**
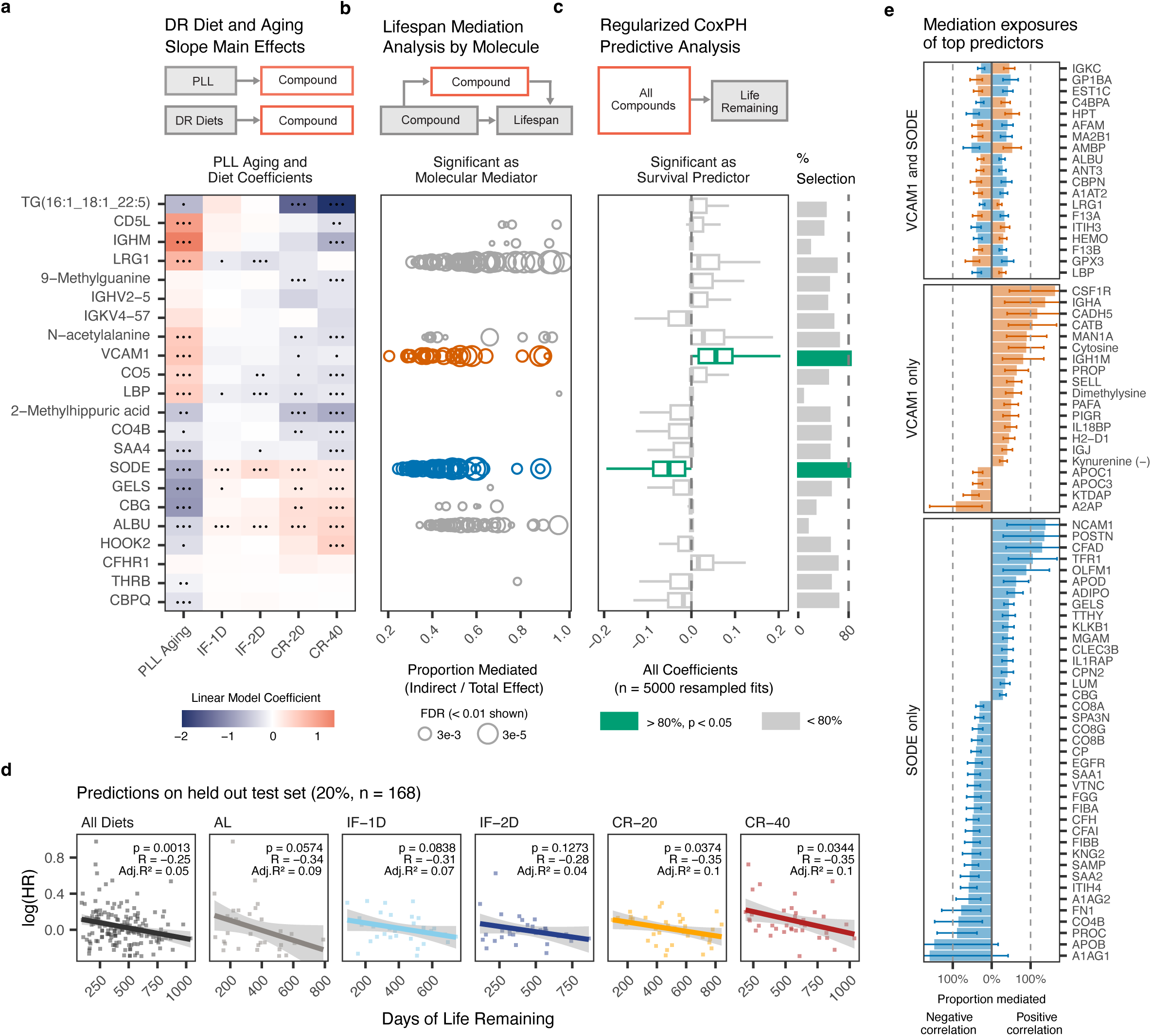
Lifespan extension is associated with both DR-dependent and independent compounds. Mediation and regularized Cox analyses use Year 2 sample data collected before the last 15% of life lived. **a,** Heatmap of PLL Aging slopes and Diet coefficients from linear mixed effects models. **b,** Dot plot of significant (FDR < 0.01) mediation relationships between compounds on lifespan, with proportion mediated < 0 or > 1 removed for clarity. Y-axis compound labels represent the mediators, and each dot represents a significant exposure. **c,** Boxplot of lasso-penalized Cox survival model shows estimates from n = 5000 resampled models, for compounds selected above a natural threshold (Ext. Fig. 3b). **d,** Predicted hazard ratios from Cox model on held out test set (20% of samples). **e,** All mediation exposures for SODE and VCAM1 with bars representing the standard error. Compounds that negatively correlate with focal mediator compounds are left of 0% line, and compounds that positively correlate with mediator compounds are right of the 0% line. •••/••/• = FDR < 0.001, 0.01, 0.05 on effect estimates.

In the regularized survival model, only SODE and VCAM1 were selected in greater than 80% of bootstrap iterations (82%, p = 0.027 and 83%, p = 0.020, respectively; Fig. 4c, Supp. Table 8), independently confirming the emergence of a DR-dependent (SODE) and DR-independent (VCAM1) lifespan signature. This indicates that the association of SODE with lifespan may be partially mediated by dietary intervention and that metabolic adaptation (via SODE) and immune aging (via VCAM1) represent parallel, independently regulated routes to longevity in this cohort. The exclusion of albumin from this more stringent selection likely reflects its covariation with other predictors in the model rather than an absence of biological relevance.

Hazard ratios from the held-out test set from the survival model were significantly correlated with remaining life (p = 0.0013, Fig. 4d). This pattern was consistent across all DR groups, with most variability explained in the CR-20 and CR-40 cohorts (R² = 10%, Fig. 4d). Among top predictors, SODE, LRG1, and VCAM1 showed the strongest pairwise correlations with remaining lifespan with Age and Diet groups (Ext. Fig. 4). The integration of distinct biological signals was further revealed in the mediation network as SODE more specifically integrated signal associated with wound healing (POSTN, complement components, fibrinogens) while VCAM1 more specifically integrated signal associated with humoral immunity (IGHA, IGH1M, IGJ, PIGR) (Fig. 4e). Further, SODE and VCAM1 both mediated the lifespan effects of ALBU and LRG1 (whereas LRG1 and ALBU did not mediate SODE or VCAM1 at the same FDR threshold), supporting the role of SODE and VCAM1 as focal downstream lifespan integrators (Fig. 4e). This distinction gives biological texture to their statistical separation and generates testable mechanistic hypotheses that are explored further in the Discussion.

### Genetic architecture of the plasma multiome links molecular variation to lifespan-associated loci

We examined genetic contributions to plasma compound abundances by quantifying QTL for each metabolite, protein, and lipid (mQTL, pQTL, and lQTL). A total of 9,599 QTL were found across all timepoints (Ext. Fig. 5, Supp. Table 9). Proteins had more traits with QTL, more QTL per trait, and were the most heritable of the three molecular classes (Fig. 5a, Supp. Table 10). Lipids exhibited the lowest median heritability and percent of traits with QTL, but showed a notable increase in heritability over time, possibly reflecting the stabilization of DR-driven lipid remodeling such that residual variation increasingly reflects stable genetic differences between individuals rather than acute dietary responses. In contrast, trait heritability decreased with age amongst proteins and metabolites (Fig. 5a). Together, these patterns suggest that the relative contributions of genetic and environmental factors to plasma molecular variation are dynamic rather than fixed and shift in a molecularly class-specific manner across the lifespan.

**Fig. 5:**
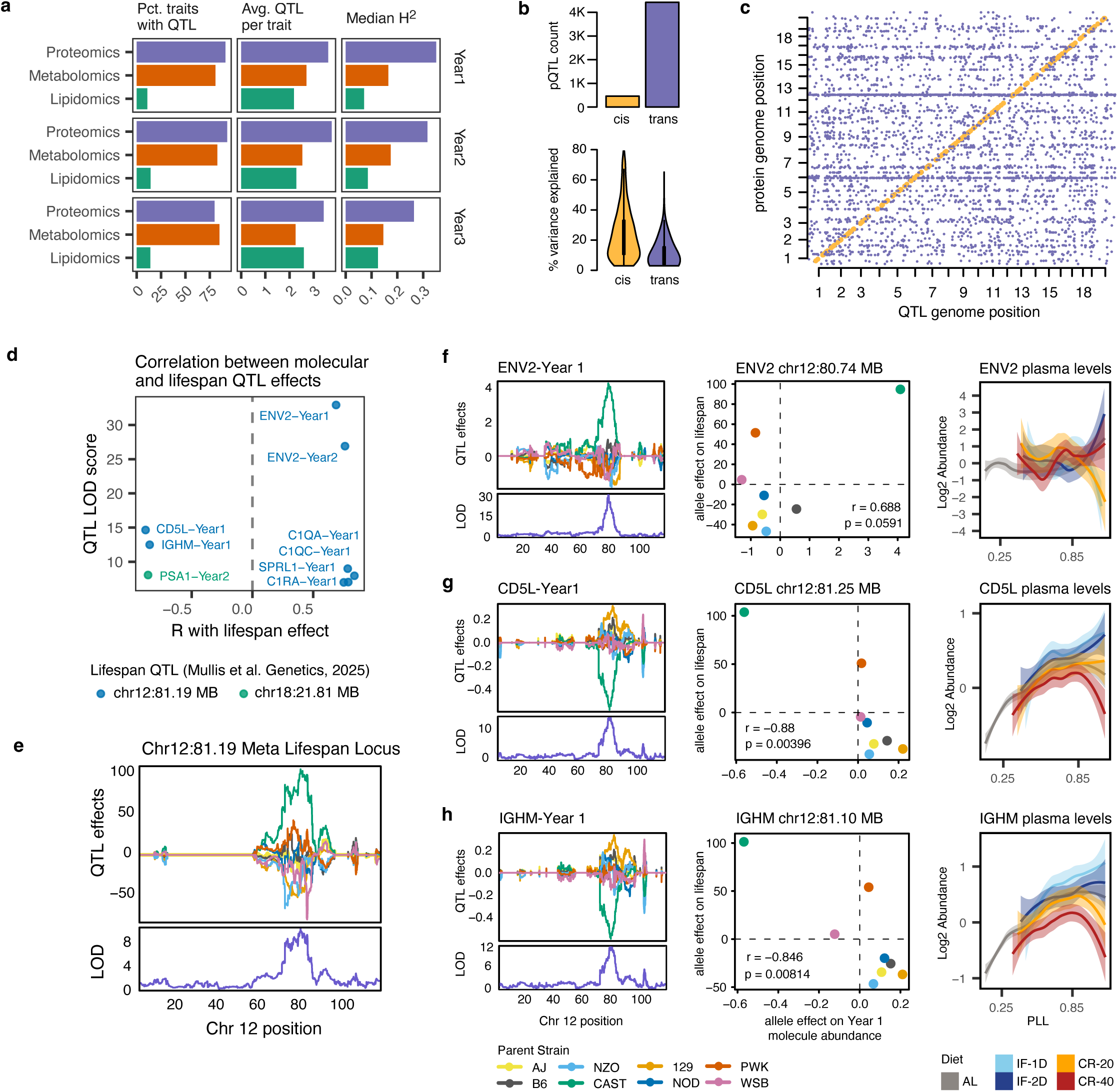
Genetic architecture of the plasma multiome links molecular variation to lifespan-associated loci. **a,** Metabolite, protein, and lipid QTL (mQTL, pQTL, and lQTL) were calculated for each compound at each timepoint. **b,** Cis-QTL were defined as pQTL falling within 1 Mb from the transcription start site (TSS) of the associated gene, and trans-QTL encompass all remaining QTL. Mapping of pQTL with cis (yellow) or trans (purple) annotation reveals genomic hotspots. **c,** Cis-QTLs describe most of the variability in the genome despite detected of more trans QTL. **d,** Six lifespan loci previously identified in a study of DO mice were overlaid with the molecular QTL resource. Nine molecular QTL shown colocalized with a 1 Mb window of lifespan and were allelically correlated with the lifespan effect. **e,** LOD and QTL effects on days of lifespan on meta lifespan-associated locus on chromosome 12. **f,** Parent allele plot of Fv4 (Year 1) at chr12:80.74 (LOD 32.9) with its allelic correlation against lifespan and pattern of change with DR over PLL. **g,** Parent allele plot of CD5L (Year 1) at chr12:81.25 (LOD 14.6). with its allelic correlation against lifespan and pattern of change with DR over PLL. **h,** Parent allele plot of Ighm (Year 1) at chr12:81.10 (LOD 12.5) with its allelic correlation against lifespan and pattern of change with DR over PLL.

We enumerated cis (pQTL falling within 1 Mb of the transcription start site for the that protein’s gene) and trans pQTL, finding 457 cis pQTL and 4423 trans pQTL, with the cis pQTL generally explaining more variance per locus than did trans pQTLs (Fig. 5b). Examination of pQTL with cis and trans annotation revealed genomic regions on chromosomes 6 and 12 with many trans pQTL (Fig. 5c). These regions have a high gene density and encode many immunoglobulin variable domains. Under closer examination, we found that the pQTL at these locations were spread over many different variable domain proteins, and showed a similar number of QTL per protein and percent of proteins with a QTL (Ext. Fig. 6).

To determine whether any of the detected molecular QTL had a direct genetic basis in longevity, we overlaid the full QTL resource with six previously identified lifespan-associated loci in DO mice^23^. This analysis revealed QTLs for 19 molecular traits that co-localize within a 1 Mb window around the lifespan locus. Of these, nine showed parent allele correlation (p < 0.1) with its lifespan effect. Four of these traits—ENV2 at year 1 and 2, IGHM at year 1, and CD5L at year 1—showed LOD scores greater than 10 near lifespan locus chr12:81.19 (Fig. 5d-e). ENV2 has a likely cis-QTL at chr12:80.74 with IGHM and CD5L having trans QTL at chr:12:81.10 and chr12:81.25 respectively (Fig. 5f-g). ENV2, IGHM, and CD5L all showed strong QTL effects driven by the CAST allele, which was the most prominent allele in the lifespan association at this locus^23^ (Fig. 5e-h). CD5L and IGHM also showed strong positive Aging slopes and moderate lifespan mediation effects (Fig. 4a-b), directly connecting this genetic locus to the molecular aging signatures described in Figure 4. Interestingly, ENV2 was highly variable with no clear pattern throughout PLL, while IGHM and CD5L increased through the Aging phase but showed a rare instance of DR-dependent variability at end-life, with the CR groups decreasing and IF and AL groups increasing during Dying phase. (Fig. 5f-h). Taken together, the co-localization of these molecular QTL with lifespan-associated loci illustrates the potential of integrating multiomic molecular profiling with genetic mapping and reinforces the importance of immune function to aging and longevity.

The remaining QTL that colocalized with lifespan locus chr12:81.19 were positively associated with the lifespan effect and were specifically classical complement cascade components (C1QA, C1QC, C1RA) and SPARC-like protein 1 (SPRL1) a glycoprotein secreted from astrocytes. In contrast, PSA1, the only trait whose QTL effects correlated with lifespan on chr18:21.81, showed a negative genetic correlation with lifespan. All correlated allelic effects were driven by the CAST allele (Ext. Fig. 7).

## Discussion

This study provides a comprehensive multiomic characterization of plasma molecular aging and dietary restriction, integrating metabolomics, lipidomics, and proteomics across the full lifespan of genetically diverse mice. Three key findings emerge. First, IF and CR converge on a shared plasma molecular program, with the magnitude of change scaling with caloric intake. Second, plasma molecular aging is biphasic, with a prolonged period of gradual change giving way to a distinct and rapidly evolving terminal phase at approximately 85% of natural lifespan. Third, two independent analytical approaches converge on SODE and VCAM1 as the plasma proteins most strongly associated with lifespan in this cohort. Together,these findings suggest that immune and lipid regulatory pathways are conserved aging and DR pathways with independent and overlapping effects, consistent with meta-analyses in humans showing that immune activity, insulin/IGF1 signaling, and lipid function are among the most consistent pathways linked to longevity^24^.

Plasma aging trajectories were biphasic, with steady rates of molecular change that sharply accelerate or decelerate at a point of terminal decline between 80% and 90% of natural lifespan. The consistency of this inflection point (Fig. 3b) across diverse data types points to a coordinated, organism-wide biological transition rather than a gradual and continuous aging process. This finding has direct parallels in human aging biology: organ-level proteomic studies in humans have suggested an aging inflection at approximately 50 years^25^, and plasma multiomic studies have identified one or more nonlinear transition points between 44 and 60 years of age^26^. In humans, the average lifespan has increased in the last 100 years, but newer studies suggest that this increase is due to a lengthening of the dying process rather than an extension of healthy life^27^. A meta-analysis of end-of-life trajectories in humans have described this period as “terminal decline”, with many findings focused on decline in the 3-7 years preceding death^28^. Similarly, imbalances in the stress-repair ratio show an inflection at age 75^29^. The convergence of mouse and human data on a biphasic model of aging underscores the importance of distinguishing healthy aging trajectories from terminal decline when identifying aging biomarkers, which is a distinction that is only possible when individual lifespan data are incorporated into the analysis.

We found terminal decline to be associated with increased levels of both proteins and metabolites implicated in nucleotide catabolic processes, which showed minimal changes during the healthy aging phase (Fig. 3f, g). This is consistent with other studies in humans that have identified increased levels of modified nucleotide components as high mortality- and disease-risk compounds, including dimethylguanosine, pseudouridine, methylguanine, methylguanosine, and methyladenosine^30–34^. Other studies have shown that nucleic acids in the plasma are biomarkers of cancer^35^ and associated with renal disease in human cohorts across multiple comorbidities^34^. We did not observe enrichment of canonical CKD biomarkers creatinine and albumin with age, but we did observe enrichment of urea-proxy homocitrulline and lipocalin-2, the latter of which has been reported as a biomarker and active protein in disease progression of CKD in mice^36^. We propose that circulating intracellular components, such as proteasome subunits and nucleosides, reflect cell death and not healthy aging. This study underscores the need for careful consideration of collection timepoint when identifying aging-related plasma biomarkers.

We found that changes in the plasma multiome due to diet are largely consistent across IF and chronic CR and scale with caloric intake, but that even more compounds change with age than with diet. While circulating lipids and amino acids have directionally correlated changes between aging and DR, proteins associated with inflammation, fibrosis, and lipid clearance and TCA cycle metabolites have counteracting effects. This finding suggests decreased beta oxidation and mitochondrial metabolism with age which can be mitigated by CR, given the significant increases in malate and 2-oxoglutarate in the CR-20 and CR-40 groups (Fig. 2b). This is consistent with human metabolomics studies showing increased TCA cycle compounds in plasma with extended lifespan^30^. Adiponectin represents a compelling candidate hormonal mediator of DR-dependent lifespan extension in this cohort given that it improves insulin sensitivity, reduces inflammation, and regulates energy metabolism^19–21^, and in this study, decreased with age and was restored by all four DR interventions.

We employed complementary mediation and survival modeling approaches to identify plasma proteins most likely to lie in longevity-associated pathways. VCAM1, an endothelial adhesion molecule whose upregulation is associated with many diseases, regulates the adherence of leukocytes to endothelial tissues in inflammatory diseases^37^ and has recently been suggested as a potential aging target^38^. A large human GWAS analysis identified circulating VCAM1 as one of four proteins detrimental to healthy aging^39^. Studies in mice have shown increases in circulating VCAM1 with aging and improvements in cognitive performance with VCAM1 deletion or anti-VCAM1 antibody treatment^40^. LRG1 is also expressed in response to inflammatory stimuli and is associated with many disease states in humans, including fibrosis, cardiovascular disease, and diabetes^41–45^. When taken in consideration with the corpus of scientific literature, our findings indicate that increased circulating VCAM1 and LRG1 reflect elevated inflammation and fibrosis with aging^46^, though VCAM1 and LRG1 levels were only modestly reduced by DR. These proteins represent DR-independent aging processes that warrant investigation as therapeutic targets.

In contrast, DR actively increased both albumin and SODE. Albumin is the most abundant protein in the plasma and has been proposed as a marker of mortality risk in humans ^47,48^. Although associations between SODE and mortality in humans are less clear, short-term DR reduces oxidative stress in humans, in some cases increasing SOD specifically^49–51^. SODE is a highly conserved protein whose expression was reported as a strong predictor of remaining lifespan in C. elegans^52^, suggesting a fundamental and conserved role in determining longevity, although this effect appears to be due to changes in transcriptional profiles and not direct effects of antioxidation. In mice, mutation of the heparin binding domain of SODE resulted in premature aging due to systemic inflammation^53^. Conversely, overexpression of SODE upregulated genes associated with energy expenditure, reducing metabolic disease under high-fat diet^54^. We posit that elevated SODE with DR may feed into other DR- and longevity-associated phenotypes in this study, including reduced inflammation and altered mitochondrial and fatty acid metabolism^55,56^. Future studies may follow up on the hypothesis that VCAM1 is implicated in humoral immune aging and SODE in metabolic adaptations with age and wound healing (Fig. 4e). From this, we propose that immune adaptations with DR affect innate rather than adaptive aging pathways, which may have important implications for understanding how DR modifies immune aging in humans.

Overlaying the molecular QTL resource with known lifespan-associated loci in DO mice^23^ revealed 19 co-localizing molecular traits, the most striking of which was the co-localization of a ENV2 cis-QTL with trans-QTL for IGHM and CD5L at chromosome 12:81.19. ENV2, a resistance gene against ecotropic murine leukemia viruses, IGHM, and CD5L all showed strong effects driven by the CAST allele, which is the same allele most prominently associated with lifespan extension at this locus^23^. Given that IGHM and CD5L both show strong positive Aging slopes and moderate lifespan mediation effects, combined with the fact that ENV2 is a cis-QTL at this locus, the lifespan-extending effect of the CAST allele at this locus may operate through ENV2-mediated suppression of viral-driven immune activation, in turn modulating circulating IGHM, CD5L, and complement activation, contributing to the lifespan association at this site.

This genetic finding independently supports the central role of immune regulation in aging and DR identified through the molecular analyses, and the full QTL resource represents a rich substrate for future investigations into the genetic architecture of aging and DR response.

## Limitations

One limitation to this study is that plasma sampling was limited to up to three timepoints per mouse. While this limitation is partially offset by converting age to proportion of life lived and the large overall sample size, future studies would benefit from increased sampling frequency. As plasma is only reflective of systemic changes rather than tissue-level changes, future studies may investigate whether transcriptomic or proteomic analyses of tissues such as liver, adipose, and immune cells can elucidate the molecular mechanisms underlying the metabolic and immune changes observed here with DR. In other DO mouse studies, transcripts associated with humoral immunity, complement factors, and VCAM1 were detected in the kidney, with complement factors also detected in the heart of aged mice^57,58^, but tissue sources for the plasma phenotypes described herein, including adiponectin and albumin, are not clear^59,60^. Other circulating proteins shown to be lifespan-associated such as GDF15 and NFL were not detected with our methodology and parallels cannot be drawn^61,62^. This study was also limited in its use of female mice only. Several studies have reported that DR effects on lifespan and the molecular responses to DR differ between sexes, with lifespan benefits in some cases reported to be larger in females than males^63^. Female mice were chosen because they are less variable and more understudied. Finally, the cause of death in this cohort was not definitively established. This ambiguity underscores the importance of interpreting the Dying phase molecular signature as reflecting end-life biology broadly rather than as any specific disease process.

## Conclusions

This study makes several contributions to the understanding of aging and dietary restriction that were not previously possible. We establish that plasma molecular aging is biphasic, with a coordinated transition to terminal decline at approximately 85% of natural lifespan, a pattern that mirrors nonlinear aging dynamics reported in humans. By specifically examining the time prior to terminal decline, we have identified VCAM1 as a marker of humoral immune aging and SODE as a marker of metabolic adaptation and wound healing. We show that DR affects metabolic and innate immune pathways but not adaptive immune pathways, a distinction with potentially important implications for understanding how DR modifies aging in humans. The multiomic QTL resource provided, and the genetic architecture of lifespan-associated plasma traits it reveals, provides a foundation for future investigations into causal drivers of aging and DR response across diverse genetic backgrounds. Ultimately, this plasma analysis permits close comparison to human studies and positions the plasma multiome as a powerful and translatable window into the biology of healthy aging.

## Supporting information

All Supplemental Tables

## Author Information

### Contributions

A.D.F., F.M., G.A.C., and B.D.B. contributed to conceptualization. J.Y.F., C.S., N.V., M.M., P.S., A.L., and B.D.B. performed data curation. J.Y.F., M.M., and B.D.B. performed formal analysis. C.S., N.V., L.J.G.C., N.O., T.N., and A.G. conducted the investigation. C.S., N.V., P.S., A.L., J.O., J.V., G.A.C., and B.D.B. contributed to methodology. C.S., L.R., and B.D.B. handled project administration. P.S. and W.L. developed the software. S.R.H., R.K., F.E.M., G.A.C., and B.D.B. provided supervision. J.Y.F. performed visualization. J.Y.F. and B.D.B. wrote the original draft. All authors reviewed and approved the final manuscript.

## Acknowledgements

We would like to thank David Botstein, Anil Raj, and Kevin Wright for helping motivate this study, Zhenghao Chen and Prateek Gundannavar for advice on data analysis methodologies and Alireza Delfarah for assistance with sample preparation.

## Data Availability

R code used for analysis and figure visualization with processed data is available at https://github.com/calico/drido-multiomic-paper. Processed data can be explored in an interactive app https://public-rstudio-connect.calicolabs.com/drido-multiomics/. QTLs from this study are available to browse at https://churchilllab.jax.org/qtlviewer/DRiDO.

## Methods

### Experiment

This study investigated the effects of various Dietary Restriction (DR) protocols on plasma proteins, metabolites, and lipids in 960 female Diversity Outbred (DO) mice. All procedures were reviewed and approved under Jackson Lab IACUC protocol 06005. The mice underwent DR intervention in multiple waves from generations 22–28 and were housed in groups of eight with only one mouse per litter. DR interventions started at 6 months of age, following randomization by cage. DR inventions included control Ad Libitum (AL), a 24-hour fast (IF-1D), and a 48-hour fast (IF-2D), 20% chronic restriction (CR-20), and 40% chronic restriction (CR-40) (Fig. 1a). All mice were fed standard chow, and the CR groups were given measured amounts at 15:00 daily to align with the beginning of the dark cycle. The 40% group was gradually restricted over the course of 4 weeks. For the CR groups, competition for food was mitigated by placing food on the floor of the cage. On Friday afternoons, a triple-ration of food was provided for the CR groups. More details about the experimental protocol, phenotyping, and collection of other physiological and functional metrics have been previously published^12^.

### Plasma Collection

Plasma was collected from retro orbital blood draws using local anesthesia (Tetracaine, proparacaine, isoflurane) in an awake mouse. 200 uL of blood were collected into an EDTA tube and placed directly on ice. Samples were centrifuged at 14000 rpm at 4C for 10 minutes. 50 uL of plasma was collected and stored at -80C. Plasma was collected at 24 (Year 1), 71 (Year 2), and 122 (Year 3) weeks of age with all animals fasted for 4 hours prior to collection. Due to mouse deaths before collection timepoints and technical outlier removal, a total of 2234 samples were analyzed (n = 940 Year 1, 823 Year 2, 471 Year 3).

### Sample Preparation

#### Proteomics

Plasma samples were prepared on the Hamilton Vantage as previously described^64^. Briefly, 2 µL of plasma were diluted in lysis buffer (75 mM NaCl, 50 mM HEPES pH 8.5, 3% SDS, with cOmplete ULTRA and PhosSTOP inhibitors). Proteins were reduced with 5 mM dithiothreitol (DTT) for 1 hour at RT, alkylated with 15 mM iodoacetamide for 1 hour at RT in the dark, and quenched with 5 mM DTT for 30 minutes at RT in the dark. Proteins were isolated using SP3 (single-pot, solid-phase-enhanced sample-preparation) beads (Cytiva). Proteins were added to beads at a ratio of 1 µg protein to 10 µg beads, then acetonitrile (ACN) was added to a final concentration of 75% and incubated for 3 hours at RT to allow protein binding. Protein-bound beads were washed twice with 70% ethanol followed by one wash with 100% ACN. The beads were resuspended in a digestion buffer (50 mM EPPS pH 8.5, 10 mM CaCl2) then digested overnight with 1 µg of Trypsin/Lys-C Mix (Promega) at 37°C, 250 rpm. Digested peptides were transferred to new plates and labeled with TMTpro 16plex Label Reagent Set (Thermo Fisher Scientific) at a 8:1 TMT:peptide ratio (w/w). The TMT labeling reaction was quenched with 1% hydroxylamine (final concentration) and incubated for 15 minutes at RT. Samples were randomized into 16-plexes such that all timepoints derived from a single individual mouse were contained within the same plex. The 126n channel was consistently reserved for the bridging sample (a bulk pool of all experimental samples). Each 16-plex was created using the combine-mix-split strategy followed by peptide cleanup using SP3 beads. Peptides were added to beads then diluted to 95% isopropanol and incubated for 20 minutes at RT for binding. Peptides were washed twice with 95% isopropanol followed by one wash with 100% ACN, then peptide-bound beads were dried under nitrogen and eluted with 5% ACN. Eluted peptides were acidified to 5% formic acid then transferred into LC-MS vials and stored at -80°C until ready for further analysis.

#### Metabolomics and Lipidomics

All LCMS-grade solvents (water, methanol, acetonitrile, IPA) were purchased from Fisher Scientific. HPLC-grade MTBE and chloroform, and LCMS-grade Ammonium Acetate were purchased from Sigma Aldrich.

##### Sample Randomization and Aliquoting

The complete sample inventory was randomized and assigned into well positions of 26 total 96 well plates such that each plate contained 6 TMT plexes with each plex containing 15 samples and 1 bridge. Randomization was done such that all samples from a single mouse were contained within one TMT plex. To avoid an additional freeze thaw, the sample vials were transferred on dry ice into their assigned 96 well format. To aliquot the plasma for extraction, 4µL of plasma was transferred from the sample vials into a 96 well plate for proteomics analysis. 30µL of plasma was transferred into 2mL glass vials for co-extraction of metabolites and lipids. Any remaining voluming in the sample tubes was pooled to create a bulk pool sample used to monitor extraction and instrument variation for metabolomics and lipidomics analyses. The bulk pool also served as the bridge sample for all TMT plexes. Samples were stored at -80C until analysis.

##### Metabolomics and Lipidomics extraction

30µL of mouse plasma was thawed on ice for 60 min prior to the lipidomics and metabolomics extraction. Metabolomics and lipidomics samples were co-extracted by LLE as previously described^65^ using a customized AL Dual Head Rail (DHR) system from Trajan Scientific equipped with glass syringe liquid handling, vortexer, centrifuge, and refrigerated vial cabinet. Briefly, 30µL of plasma was aliquoted into 2mL glass vials and 790µL of -20C (40:37, v/v) methanol/water containing internal standards listed in the Table 1 was added to the vial and vortexed. Two lipid extractions of 800µL MTBE were performed and pooled to create the lipid extract. The remaining aqueous phase was removed for the metabolite extract. Both extracts were dried under nitrogen gas while maintained at 4C. The lipid fraction was reconstituted in 750 µL of 4:2:1 IPA:MeOH:Chloroform + 5.5mM ammonium acetate containing one internal standard for every quantified lipid class. The metabolite fraction was reconstituted in 400 µL of 4:5:1 ACN:MeOH:Water including the post-extraction standards listed in Table 1.

### Data Acquisition

#### Proteomics

Peptides were analyzed on an Orbitrap Eclipse mass spectrometer coupled to an UltiMate 3000 HPLC (Thermo Fisher Scientific). Peptides were separated on an IonOpticks Aurora Ultimate C18 microcapillary column (75 µm inner diameter x 25 cm length, 1.7 µm particle size, 120Å pore size) operating at 60°C with a flow rate of 300 nL per minute. The total LC-MS run length for each sample was 185 min including a 165-min gradient from 6% to 35% acetonitrile in 0.1% formic acid. The data was collected in data-dependent acquisition (DDA) mode. Four different FAIMS Pro compensation voltages (-40, -50, -60, -70 V) were used, each with a 1.25 s cycle time. High-resolution MS1 scans were acquired in the Orbitrap mass analyzer with a resolution of 120,000 and scan range of 400-1600 m/z. The automatic gain control (AGC) target was set to standard with ‘Auto’ max injection time and 30% RF lens value. Dynamic exclusion was enabled after 1 time with a 60 s duration and a 10 ppm mass tolerance. For MS2 spectra, ions were isolated with the quadrupole mass filter using a 0.7 m/z isolation window. The MS2 product ion population was analyzed in the quadrupole ion trap (CID, 1 × 104 AGC, 35% normalized collision energy, and 35 ms max injection time). The MS3 scan was analyzed in the Orbitrap (HCD, 50,000 resolution, 1 × 105 AGC, 200 ms max injection time, 45% normalized collision energy) and synchronous precursor selection (SPS) precursor ranges were set to 455-1600 m/z. Up to ten fragment ions from each MS2 spectrum were selected for MS3 analysis using SPS. The mouse UniProt proteome (proteome ID UP000000589) was used for real time search and a maximum of 2 missed cleavages and 1 differential modification was allowed. The maximum search time was set to 35 ms and a cross-correlation score of 1, delta cross-correlation score of 0.1 and precursor of 10 ppm for charge states 2, 3 and 4 was used.

LC-MS data were processed using a previously described software pipeline^66^. Raw files were converted to mzXML format and searched against a mouse UniProt database using the SEQUEST algorithm. MS/MS spectra were matched with fully tryptic peptides using a precursor mass tolerance of 20 ppm and fragment ion tolerance of 0. Carbamidomethylation of cysteines (+57.021 Da) and TMT modification of peptide N-termini and lysine residues (+304.207 Da) were set as static modifications. Oxidation of methionine (+15.995 Da) was set as a differential modification. Peptide matches were filtered to a 1% false discovery rate (FDR) using linear discriminant analysis (LDA) as previously described (Huttlin et al. 2010). Non-unique peptides that matched to multiple proteins were assigned to proteins that contained the largest number of matched redundant peptide sequences using the principle of Occam’s razor (Elias and Gygi 2007). TMT reporter ion quantification was performed by extracting the most intense ion within a 0.003 m/z window at the predicted m/z value for each reporter ion.

#### Lipidomics

Lipidomics data was acquired on a Thermo ID-X Tribrid mass spectrometer connected to an Advion Triversa Nanomate equipped with an electrospray chip containing a 20x20 grid of 5µm internal diameter nozzle tips, allowing direct infusion at flow rates between 100-500 nL/min. Negative mode acquisition was used to acquire phospholipids, fatty acids, ceramides, and sphingolipids. Positive mode acquired phospholipids, glycerolipids, MGs, ceramides, cholesterol, cholesterol esters, sphingolipids, and carnitines. The direct infusion method operated as follows: first, MS^1^ full scans were collected for 0.2 min to allow the electrospray to equilibrate. This data was discarded. Second, SIM scans were collected in 100 m/z windows between 200 m/z - 1200 m/z with 25 m/z overlapping between windows. A second set of SIM scans were collected between 700 m/z - 900 m/z using 20 m/z windows and 15 m/z overlapping between windows. Third, 482 combined MS^2^ scans of 1 m/z width were collected and centered at the mass of lipids previously determined to be detectable in a sample pool. Additionally in positive mode, Triacylglycerol (TG) quantification was collected via MS^3^ using CID fragmentation and measured the linear ion trap from ammoniated precursors. The set of all MS^3^ scans to monitor for TGs were defined via an initial separate analysis of a sample pool, where based upon the observed precursor mass and observed MS/MS fragments, all possible MS^3^ precursors originating from were enumerated. The total run time was 7 minutes in negative mode and 5.7 minutes in positive mode, generating at least 3 replicate measures for each scan. Analyses which showed low MS signal, were rerun using a replicate plate.

Lipid identification was performed via two distinct processes, one for MS3 scans data (applied only to TGs), and the other for MS1 + MS2 scans data (applied to all other lipids).

Using the predefined MS^3^ scans, all three acylium ion (indicative of a single acyl chain) were measured for a given TG, and quantified via the sum of the intensities of all of its acyl chains. TG quantities were then normalized by dividing by the summed acyl chain intensities of an internal standard TG. Due to the ion isolation scheme, each measured MS^3^ fragment ion was derived from a single TG species.

For all other lipids, a combined MS^1^ + MS^2^ search was used, First, all bulk pool samples were searched against an *in silico* library^67^ with theoretical fragments tagged when diagnostic of specific lipid classes,or indicative of the presence of specific acyl chains (sn1, sn2, sn3, etc). A representative adduct was defined for each lipid class, and a series of strict lipid class-specific search parameters were applied, requiring significant agreement in spectral matching (see habc_v4_bulkpool_20250116.json in the associated R package). Using the compounds matched by this criteria, a subset of the *in silico* library was created containing all adduct forms of matching compounds. A looser set of “biological parameters” were defined for subsequent matches (see habc_v4_biological_neg.json and habc_v4_biological_pos.json), which was then applied to all biological samples. In order to account for isobaric lipid species and fragments, quantities were split based on unique information (when possible), and if no unique information was possible, an approximate “split ambiguous fragments” step was carried out, whereby MS^2^ speaks were split based on the relative abundance of respective precursor ions. Values were normalized to a corresponding lipid class internal standard species, and further normalized within each plate.

The analysis pipeline is provided in ‘docr_lipids.R’, ‘habc_lipids.R’, and ‘habc4_lipids.R’ in this manuscript’s associate R package, with a complete implementation of the pipeline available in the open-source R package mzkitcpp (https://github.com/calico/mzkitcpp)

#### Metabolomics

Metabolomics samples were centrifuged at 18,000 g for 5 min at 4C in the 96 well plate, and four replicates of 80μL supernatant was moved to a new 96 well plates. Analysis of polar metabolites was performed on a Vanquish HPLC coupled to a Q Exactive Plus mass spectrometer (Thermo Fisher Scientific). Metabolites were analyzed in negative ionization mode with a SeQuant® ZIC-pHILIC column (5 μm, 200 Å, 150 × 2.1 mm) where the mobile phase A was 20 mM ammonium carbonate in water (pH 9.2) with 1 μM medronic acid (Agilent, USA), and mobile phase B was 95:5 (v/v) acetonitrile/mobile phase A at a flow rate of 150 μL/min and the gradient was t = −6, 84.2% B; t = 0, 84.2% B; t = 4, 84.2% B; t = 10, 47.4% B, t = 13, 15.8% B; t = 16, 15.8% B; t = 17, 84.2% B; t = 19; 84.2% B. Data were acquired using data-dependent acquisition (DDA) mode with the following parameters: resolution = 70,000, AGC target = 3 × 10^6^, maximum IT (ms) = 100, scan range = 65-885. The MS2 parameters were as follows: resolution = 17,500, AGC target = 1 × 10^5^, maximum IT (ms) = 50, loop count = 6, isolation window (m/z) = 1, (N)CE = 20, 50, 100; Apex trigger(s) = 3–15, dynamic exclusion(s) = 15.

Metabolites were analyzed in positive ionization mode using a Agilent InfinityLab Poroshell 120 HILIC-Z (2.1 × 150 mm, 2.7 µm, PEEK-lined) where the mobile phase A was 10 mM ammonium formate, 0.1% formic acid, 1uM medronic acid in 100% LCMS grade water, and mobile phase B was 10 mM ammonium formate, 0.1% formic acid, 1uM medronic acid in 90:10 acetonitrile:water at a flow rate of 350 μL/min and the gradient was t = −3, 98% B; t = 0, 98% B; t = 3, 98% B; t = 9, 85% B, t = 11.2, 60% B; t = 11.9, 5% B; t = 13.3, 5% B; t = 14, 98% B; t = 15; 98% B. Data were acquired using data-dependent acquisition (DDA) mode with the following parameters: resolution = 70,000, AGC target = 3 × 10^6^, maximum IT (ms) = 100, scan range = 65-885. The MS2 parameters were as follows: resolution = 17,500, AGC target = 5 × 10^5^, maximum IT (ms) = 100, loop count = 4, isolation window (m/z) = 1, (N)CE = 20, 50, 100; Apex trigger(s) = 3–15, dynamic exclusion(s) = 7. Analyses which showed low MS signal were rerun using a replicate sample plate.

For metabolomics annotation, raw files were converted to mzML using Proteowizard Version 3 (https://proteowizard.sourceforge.io). Compound identification and peak grouping was performed using the OpenCLaM R package (https://github.com/calico/open_clam). Peaks were matched within 10 ppm for the precursor mass and 20 ppm for fragment masses. Fragmentation and retention times were compared to an in-house library generated from authentic standards for metabolomics. Following automated annotation, a subset of data was manually inspected, validated, some identities were reassigned, and some peak integrations manually corrected using MAVEN2^67^ (https://github.com/eugenemel/maven). Based on the retention time and accurate mass, compound identifications were assigned for the whole dataset. Peak intensities were quantified by calculating the sum of the peak maximum intensity along with the scans immediately preceding and following that maximum (MAVEN “peak area top”). The value for each peak was log_2_ transformed for analysis.

### Data Availability

Metabolomics and lipidomics data are available on Metabolomics Workbench^68^, Study ID ST004829.

### Data Normalization

#### Proteomics

Proteomics data was normalized using the multiplexed mass spectrometry data analysis framework msTrawler^69^. Peptide scans with low signal (signal-to-noise (SSN) ratio sums below 20 or less than 2 non-zero intensities) were removed. Individual intensity values were divided by the sum of intensities for the scan, and intensities with ratios less than 1% were flagged as below the limit of detection (LOD). Scans with less than 4 values over the LOD were removed. Remaining values below the LOD were imputed using a Gaussian distribution around the LOD of the aggregate SSN ratio sum. All remaining values are log_2_ transformed and mean-centered per scan. Outlier values are detected by fitting a channel-specific and an intercept-only model with intensity values with higher SSN weighed more heavily than less reliable measurements. Scans with absolute residual values exceeding 3 for either model are removed. All peptide scans annotated for a specific protein were summed to generate a single value for each protein. Normalization factors were calculated using scans with standard deviations below the median. For each channel in each plex, the mean log_2_-transformed intensity was calculated, followed by a grand mean across all plexes. The normalization factor (channel mean minus grand mean) was then added to the corresponding channel intensities. Plexes were harmonized by subtracting the bridge value for each plex and protein from the biological samples to zero-center the data over the plex values. Missing values were not imputed.

#### Lipidomics

Raw lipid ion intensities (MS^1^ and MS^3^) were first normalized by dividing each lipid by class-wise Internal Standards (IS) to correct for global signal variability. IS-normalized lipid intensities were then divided by their corresponding BulkPool median to batch-correct and center the data. Samples ionized in positive mode with greater than 80% missing data are excluded, and samples ionized in negative mode with greater than 50% missing data are excluded. Triacylglycerols (TGs) detected in less than 75% of the remaining biological samples are removed, and other lipids detected in less than 60% of the remaining samples are removed. The IS-normalized, BulkPool-centered intensities were log_2_-transformed. Remaining missing values were imputed by random sampling over the LOD of the lipid-adduct group with a standard deviation of 0.15, where the LOD was calculated as the half-minimum value of the lowest detected value for each lipid-adduct feature. For each feature on each plate, systemic signal drift and well-dependent bias was reduced by regressing out the effects of injection order and well position from the final dataset.

#### Metabolomics

The data normalization procedure was performed separately on positive and negative ionization mode data before combining the datasets. Original samples that were re-run due to poor signal intensity were removed. Compounds with manually identified split chromatography peaks were summed. For each feature on each plate, systemic signal drift was reduced by regressing out the effects of injection order from the dataset. Regressing out injection-order dependent effects centered the data over zero for each compound and removed batch differences between plates. Outliers were those determined to be more than 3 standard deviations above or below the mean for the first three principle components from PCA analysis (R *prcomp()*) and were removed. Sample-to-sample intensity differences were reduced by using a Probabilistic Quotient Normalization approach, in which each sample is divided by its median feature difference against a reference^70^. For each feature on each plate, plate position dependent effects were identified and reduced by regressing out the effect of well position from the final dataset. Missing values (4.2% of the data) were not imputed in the final dataset for statistical analyses. Positive and negative ionization mode data was combined and treated as one dataset for statistical analyses.

### Statistics

#### FDR

Unless stated otherwise, all FDR thresholds were assigned using Storey’s q-value^71^ (R *qvalue*) grouped by data acquisition platform.

#### Covariates

All models excluding the regularization analysis (below) examined main effects for diet and age (as timepoint or proportion of life lived) with bodyweight and sample collection day covariates. Bodyweight was a numeric term as the total bodyweight of mice at the time of sampling in grams, and reduced modeling error per lower AIC across all data types (Ext. Fig. 1). Coefficient values for bodyweight terms between models were correlated. Sample day collection was a categorical term that identified whether an animal in the CR-20 or CR-40 groups had plasma collected on Monday versus any other day of the week. Monday sample collections for these animals were marked by an extended fasting period due to all weekend food being provided on Friday for CR groups. This led to a longer fasting period prior to the blood draw, as the CR animals consumed all their food before the weekend was over (Fig. 1a). Inclusion of the fasting term in the model reduced error in the metabolomics data, and had a negligible impact on the proteomics and lipidomics data (Ext. Fig. 1). For consistency, the model containing fasting was used across all data types.

#### Linear Mixed Effects Model

The fixed diet effects were determined using the entire dataset using a linear mixed effects model (R *lme4*) and with derived p-values (R *lmerTest*) (*∼ diet + age + bodyweight + fasting + (1|mouse_id)*). Percent life lived (PLL) linear slopes were determined on Year 2 and Year 3 samples stratified by sample collection before or after PLL = 0.85 (*∼ diet + PLL + bodyweight + fasting + (1|mouse_cohort)*. In heatmaps, if metabolites detected in both modes were selected for display, the mean coefficient and maximum pvalue between the two is shown. For selection of top Diet and Age compounds, triglycerides are removed, due to their consistent and high signal effect. Coefficients above 2 or below -2 are shown at maximum value.

#### Generalized Additive Models

Non-linear trajectories of PLL were determined on Year 2 and Year 3 samples using generalized additive models (GAM) (R *mgcv*) with default parameters. Smooth terms were calculated for each trait as a function of PLL (*∼ diet + s(PLL) + bodyweight + fasting + (1|mouse_id)*). FDRs were calculated using Benjamini Hochberg. Traits with FDR < 0.05 for s(PLL) and R^2^ > 28% were selected for visualization and trajectory curvature calculations, as R^2^ = 28% was the upper threshold for which metabolomics internal standards showed association with PLL. Significant predicted PLL values were de-dimensionalized using PCA (R *prcomp()*). The number of clusters to use was determined using a gap statistic (R *cluster::clusGap(),* optimal k = 10). Cluster identities were assigned using weighted K-means clustering on dimensionality reduced data with PCA variance used for weighting (R *flexclust::kcca()*, 10 dimensions) (Ext. Fig. 2a). The point of maximum curvature (PMC) was calculated from the absolute maximum of the second derivative of the GAM fit. The median PMC across all data was calculated at 0.85 PLL from 817 models with linear slopes (PMC = 1) removed (Fig. 3b).

#### GO Biological Process Enrichment

For pathway enrichment of proteins, protein names were converted to gene names and matched to EntrezID with R *AnnotationDbi.* PLL *Aging* and *Dying* models for proteins were filtered to compounds with FDR < 0.01, and enrichment was performed with R *clusterProfiler::enricher()* for each of the significant *Aging* and *Dying* groups, separated by compounds that were depleted versus enriched. The full list of annotated proteins detected in the dataset was used as the background list. Protein GO enrichment was filtered to pathways from the Biological Processes category.

For pathway enrichment of metabolites, compounds were matched to their InChiID and filtered into four groups using the same parameters as for proteins. Each group was enriched for pathways using a fold and enrichment and hypergeometric test from Inchi-assigned pathways from IDSL.GAO^72^: https://goa.idsl.site/goa/

For both proteins and metabolites, pathways were filtered to those with 4 or more compounds, and significant pathways containing identical lists of traits were reduced to the single pathway with the highest specificity. Fold enrichments for depleted pathways are sign flipped for visualization. Redundant pathways were further reduced by discarding any term with Jaccard similarity ≥ 0.5 and keeping the term with higher fold enrichment. Any significant (FDR < 0.01) compound for PLL *Aging* or *Dying* linear slope was used, independent of whether or not it was assigned a significant non-linear GAM fit.

#### Mediation Analysis

The mediation analysis used a three-model mediation framework to investigate relationships affecting lifespan. Analyses were restricted to year 2 samples with a PLL < 0.85. The first exposure–mediator approach uses DR as the exposure with individual molecular abundances as mediators (Ext. Fig. 3a). The second approach uses individual molecular abundance traits as exposures, with a second molecular trait as the mediator (Fig. 4b). All models were adjusted for body weight, fasting, and mouse cohort. The molecule-molecule approach was additionally adjusted for diet. For a given exposure X, mediator M, outcome Y (lifespan), and covariates C:

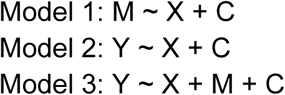

The coefficient of X in Model 1 is denoted a; the coefficient of X in Model 2 is the total effect c; the coefficient of X in Model 3 is the direct effect c′; and the coefficient of M in Model 3 is denoted b. The indirect (mediated) effect was estimated as the product a*b, with statistical significance determined using the Sobel test.

#### Lasso-penalized Cox survival model

The regularization analysis was restricted to year 2 samples with a PLL < 0.85. Batch effects from diet, fasting, and scaled bodyweight were regressed out of the data prior to analysis (*limma::removeBatchEffect()*). Data were stratified by DR intervention group, fasting group, and lifespan quartile and split into training (80%) and test (20%) sets. A LASSO-penalized Cox proportional hazards model was fit with R *glmnet::cv.glmnet()*) with α = 1, with the penalty parameter λ selected as the value minimizing cross-validated partial likelihood deviance (λ.min). Feature stability was assessed by bootstrapping the training set 5,000 times with replacement and refitting the cross validating each iteration. Per-feature selection frequency and mean coefficient were the primary metrics for further analysis (Ext. Fig. 3b). P-values were obtained by fitting an unpenalized Cox model with LASSO-selected features as covariates.

### Genetics

#### Genotyping

The mouse universal genotyping array (GigaMUGA; 143,259 markers)^73^ was used to obtain genotypes for all mice in the study. DNA was extracted and genotyped from tail tips by Neogen Genomics (Lincoln, NE, USA). Founder genotypes were constructed using the “qtl2” package in R^74^. Samples with call rates at or above 90% were retained for analysis. GCRm39 assembly genome coordinates were used, with gene locations retrieved from Mouse Genome Informatics databases^75^.

#### Additive whole genome scans

Genetic analysis was conducted using the “rqtl2” package in R^74^. Whole-genome scans were performed for each molecular phenotype using the *scan1()* function utilizing a mixed effects model in which a focal trait is regressed on 8-state allele probabilities for each individual. For each molecule, normalized abundance at each timepoint (yearly measurement) was treated as a separate phenotype. Any molecular phenotypes with measurements in fewer than 100 animals were excluded from genetic analysis. For year one phenotypes, DO generation wave and the weekday on which samples were collected were included in the mixed effects model as fixed effects; here, diet was excluded from the model as year one samples were collected prior to the administration of dietary interventions. For year two and three phenotypes, DO generation wave, weekday of collection, diet, and an interaction term between weekday and diet were included as fixed effects. In all mixed effects models, kinship matrices derived from the “rqtl2” function *calc_kinship()* using a “leave one chromosome out” scheme were included as a random effect. QTL were called using the “rqtl2” function *find_peaks()* with the following settings: ‘drop’ = 2, ‘peakdrop’ = 3, and ‘threshold’ set to one of the two statistical thresholds described below. Approximate confidence intervals around each QTL were defined using a 2LOD interval around the sentinel marker, as previously described^76,77^.

For each molecular trait, an individual significance threshold of ⍺ = 0.05 was established using 1,000 unrestricted permutations of the data, with permutations consisting of a whole-genome scan in which molecular abundance values were randomized across all individuals. The maximum LOD score observed in each permutation was recorded, resulting in a distribution of 1,000 maximum LOD scores. The 95^th^ percentile of this distribution was used as the significance threshold (⍺ = 0.05) for the corresponding meta-trait. QTL with LOD scores greater than this threshold were considered significant at our permutation-based threshold. In addition to this significance level, a significance level of LOD ≥ 6 was also used to identify loci contributing to variance in molecular abundance.

#### Heritability estimation

Heritability of each yearly molecular trait was performed via unweighted residual maximum likelihood (REML) in “rqtl2” using the additive covariates described above. The standard error of each heritability estimate was estimated via jackknifing.

#### Effect size estimation and percent variance explained

Best linear unbiased predictors (BLUPs) were computed for all QTL using the *scan1blups()* function in “rqtl2” on yearly molecular abundance values using the additive covariates described above. For each QTL, the fraction of phenotypic variance explained was calculated using the LOD score at the peak genotyping marker via the formula^78^:

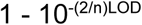

Where n is the sample size (i.e., animals) and LOD is the LOD score of the selected marker.

#### Variant association/fine mapping

Variant association mapping was conducted within a 2LOD interval around the sentinel marker of each QTL. Variant association was performed via the *scan1snps()* function in “rqtl2” using the same additive covariates detailed in the Methods section “Additive whole-genome scans” above. Variant and gene SQLite datasets used in this analysis are available at the “rqtl2” user guide website: https://kbroman.org/qtl2/assets/vignettes/user_guide.html.

#### Colocalization

For lifespan, co-localized molecular QTL were those defined as having a QTL position within 1 MB of the pre-identified lifespan QTL^77^ with a QTL confidence interval of < 10 MB within which the lifespan QTL fell. For each colocalized QTL, we compared the allelic effects at the sentinel marker on both molecular abundance and on lifespan using BLUPs from Mullis et al. 2025^77^. Pearson correlation coefficients and corresponding p values were calculated using R *cor.test()*.

---

**Extended Fig. 1:**
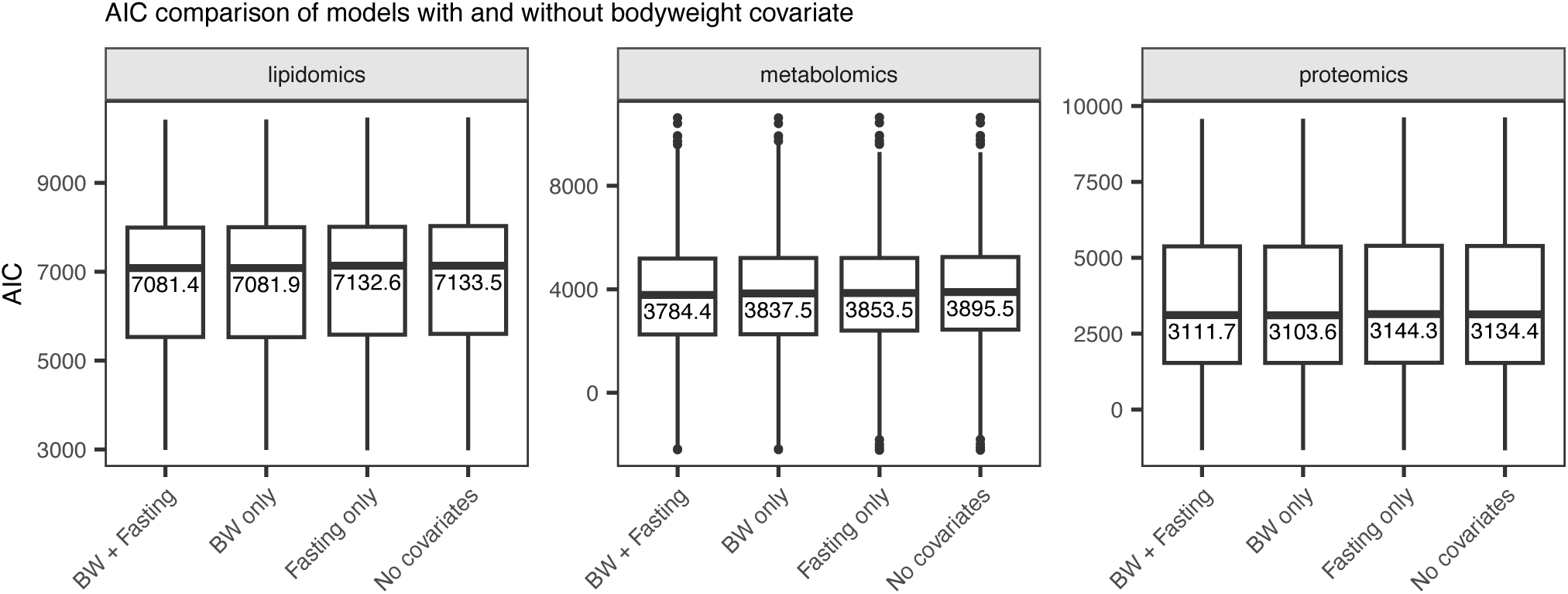
Validation of fasting and bodyweight terms in models. Exploratory data analysis found that there was a significant effect of the day of the week of the blood draw on both the metabolome and lipidome. Blood draws on Mondays for the 20% and 40% CR groups showed substantial changes relative to other samples, presumably due to triple feeding on Fridays leading to a longer fast before blood draws on that day (Fig. 1a). AIC comparison of linear mixed effects models comparing inclusion or exclusion of bodyweight and fasting covariate indicate a lower prediction error with bodyweight for all data types, and with fasting for metabolomics. For consistency across all data types, both covariates were included for main effects analysis.

**Extended Fig. 2.**
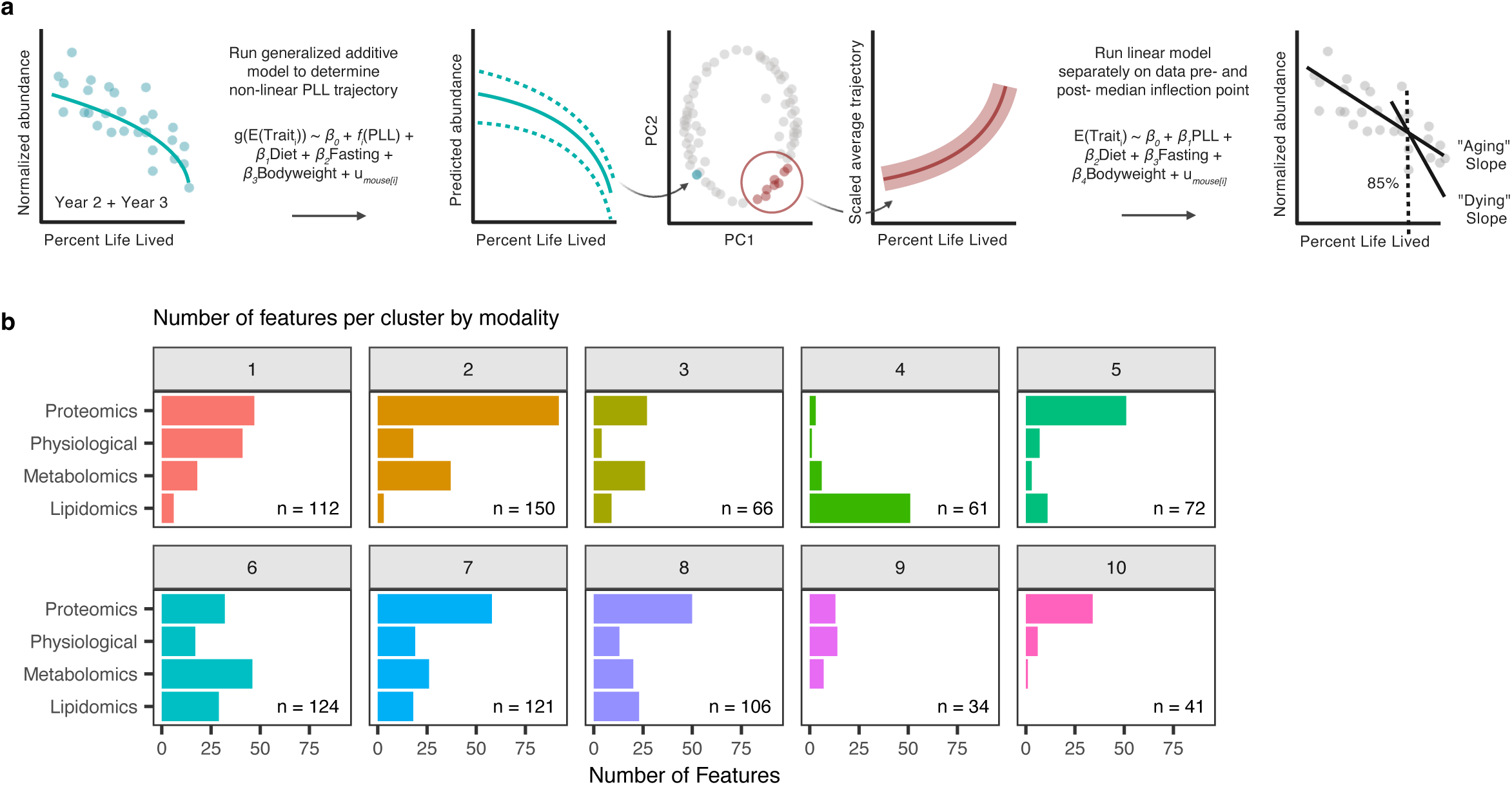
Non-linear PLL generalized additive model analysis. **a,** Modeling schematic for the quantification of non-linear aging trajectories and linear healthy aging (Aging) and terminal decline (Dying) slopes over the median maximum curvature point. **b,** Non-linear proportion of life lived (PLL) trajectories were clustered by trajectory pattern. Clusters were representative of various data types, illustrating consistency of effect versus the capturing of a data-type specific signal.

**Extended Fig. 3.**
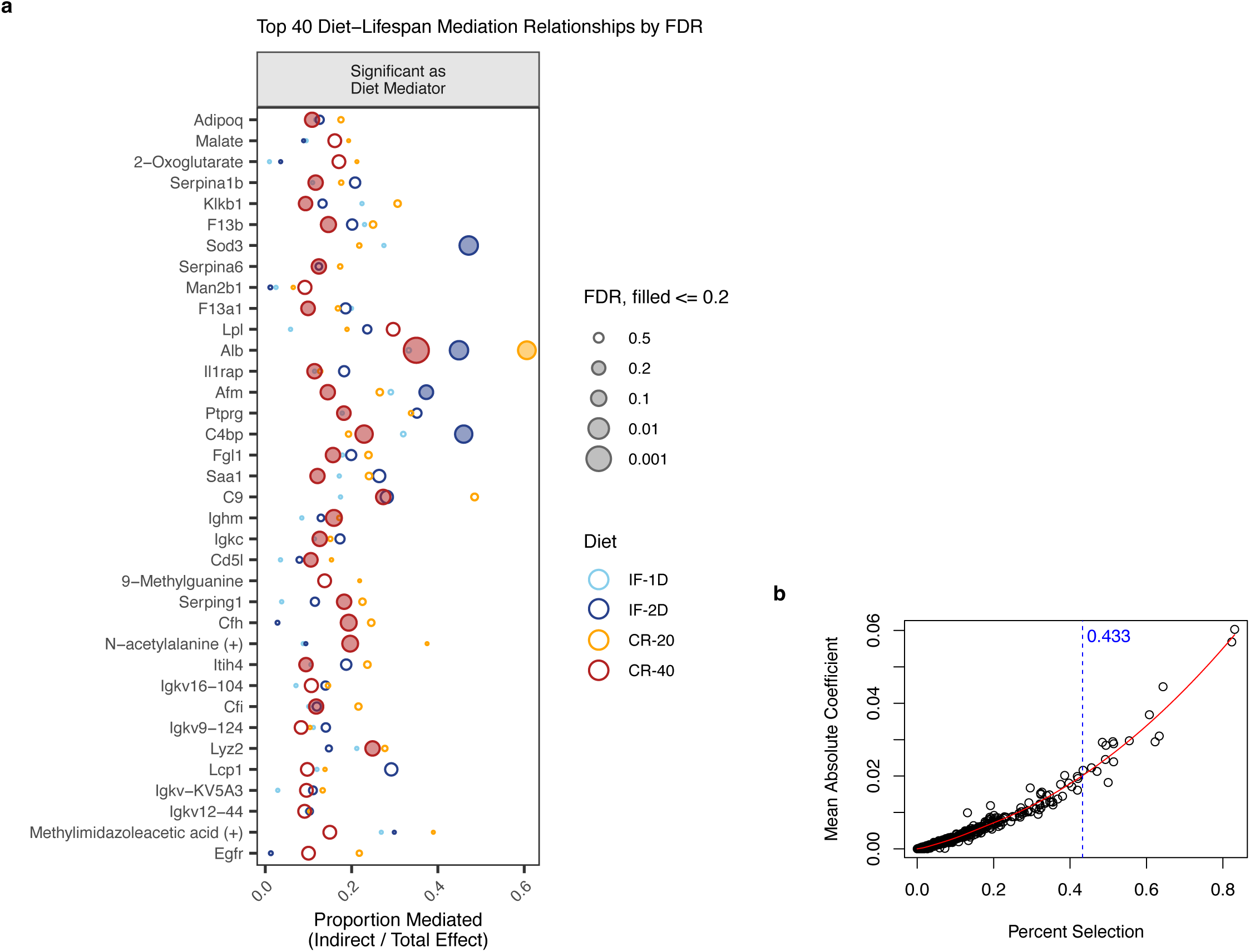
Diet mediation model and lasso selection. **a,** The top 40 molecular mediators of lifespan by DR intervention were selected by lowest FDR. Only Albumin (Alb) had an FDR < 0.01 for CR-40. This analysis use Year 2 sample data collected before the last 15% of life lived. Color represents diet, size represents FDR, and the x-axis is the proportion of DR-lifespan relationship that is mediated by the molecule on the y-axis. **b,** The number of predictors to visualize in Fig. 4c from the Lasso CoxPh model were chosen above the elbow point of the data modeled by mean absolute coefficient vs. percent selection over n = 5000 models.

**Extended Fig. 4:**
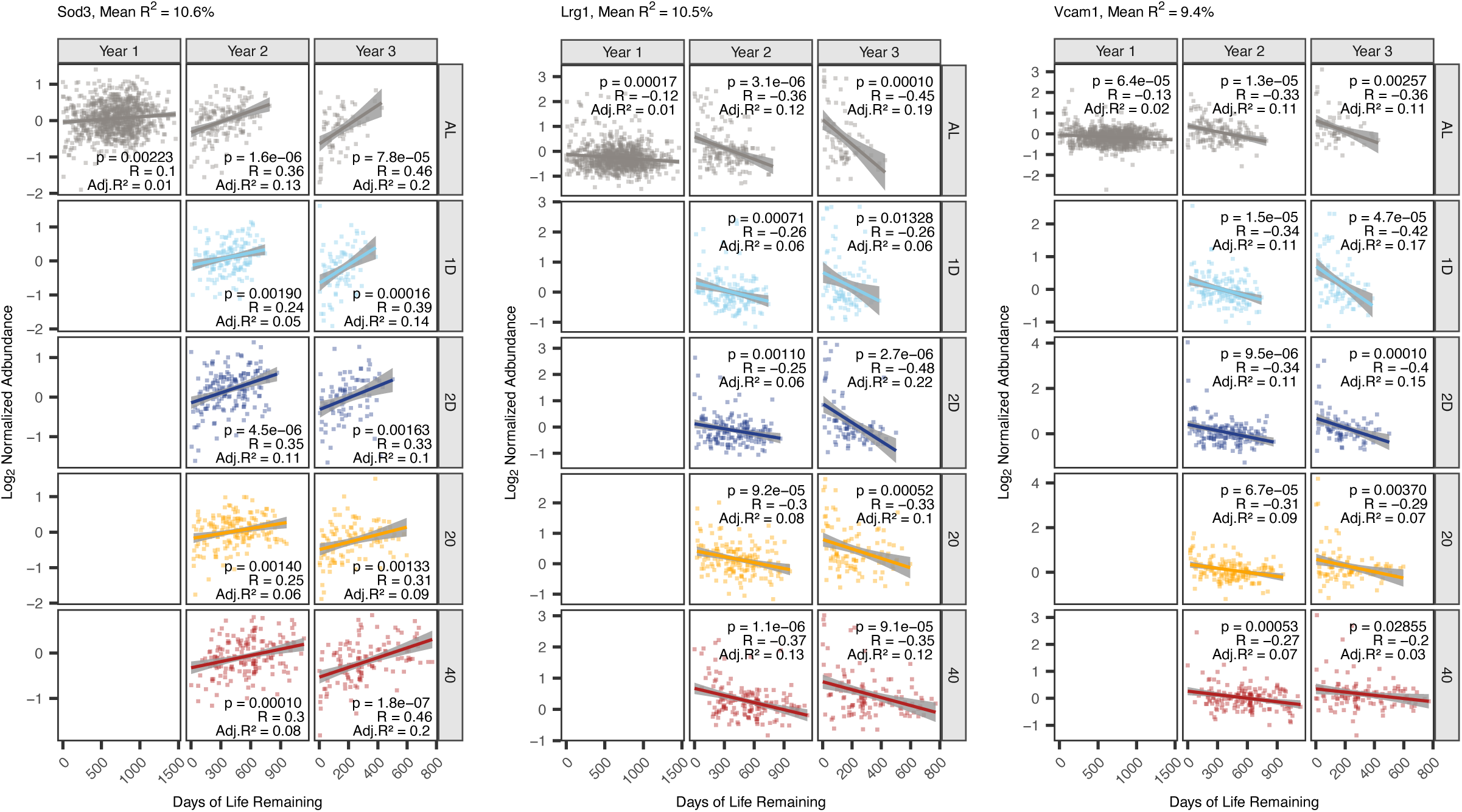
Correlations on days of life remaining and normalized abundance of top predictive plasma compounds. Superoxide dismutase (SODE), leucine-rich alpha-2 glycoprotein 1(LRG1), and vascular cell adhesion molecule 1 (VCAM1) showed the greatest correlations with days of life remaining amongst lasso-selected compounds. Normalized log2 abundance values are compared via pairwise correlation to days of life remaining for each sampling timepoint and diet intervention group.

**Extended Fig. 5:**
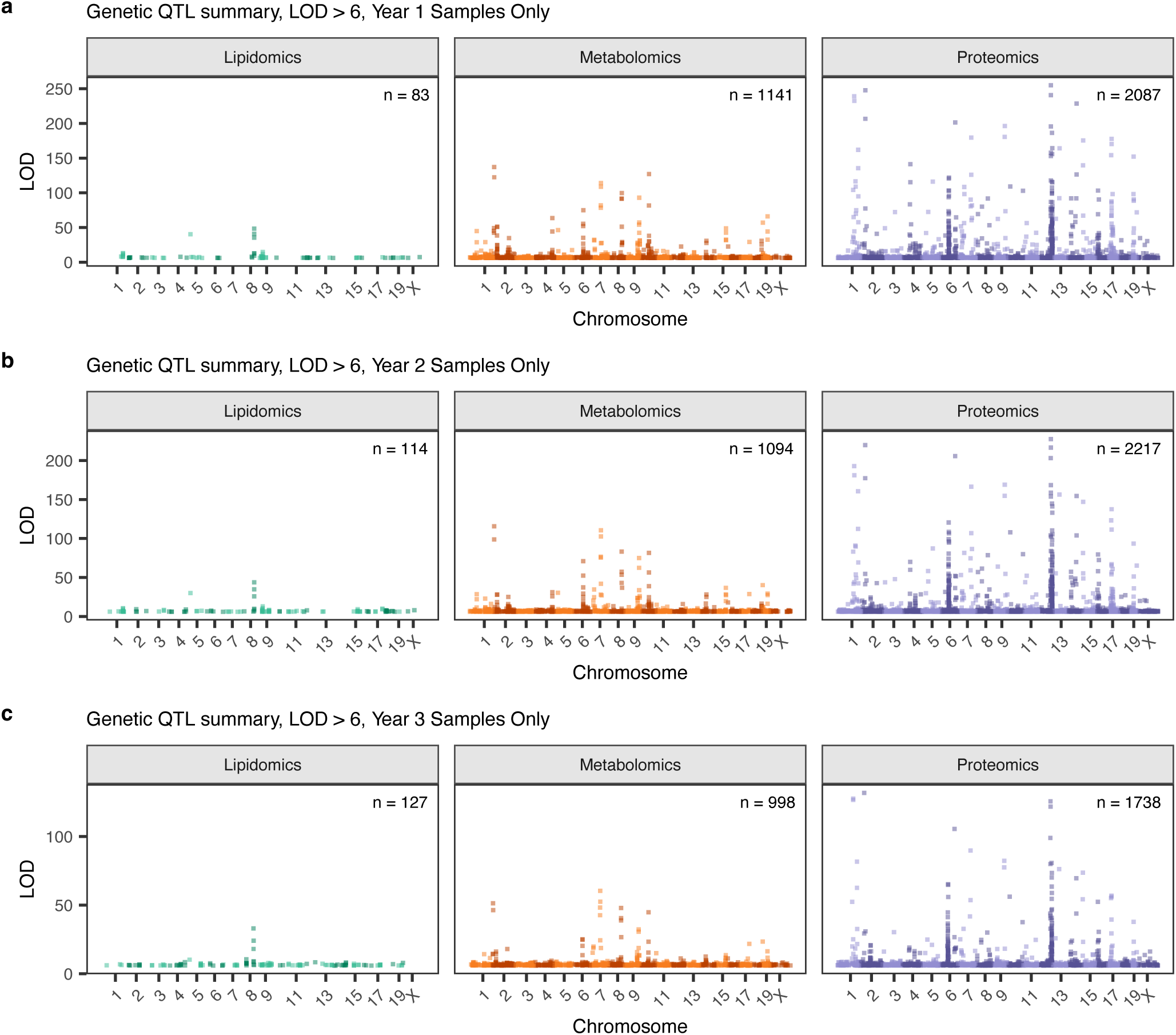
Manhattan plots of all data. Genetic QTL analysis on all plasma samples per year, accounting for DR group, cohort, and kinship, resulted in the detection 9599 QTL across lipids, metabolites, and proteins for (**a**) Year 1, (**b**) Year 2, and (**c**) Year 3 (Supp. Table 9).

**Extended Fig. 6.**
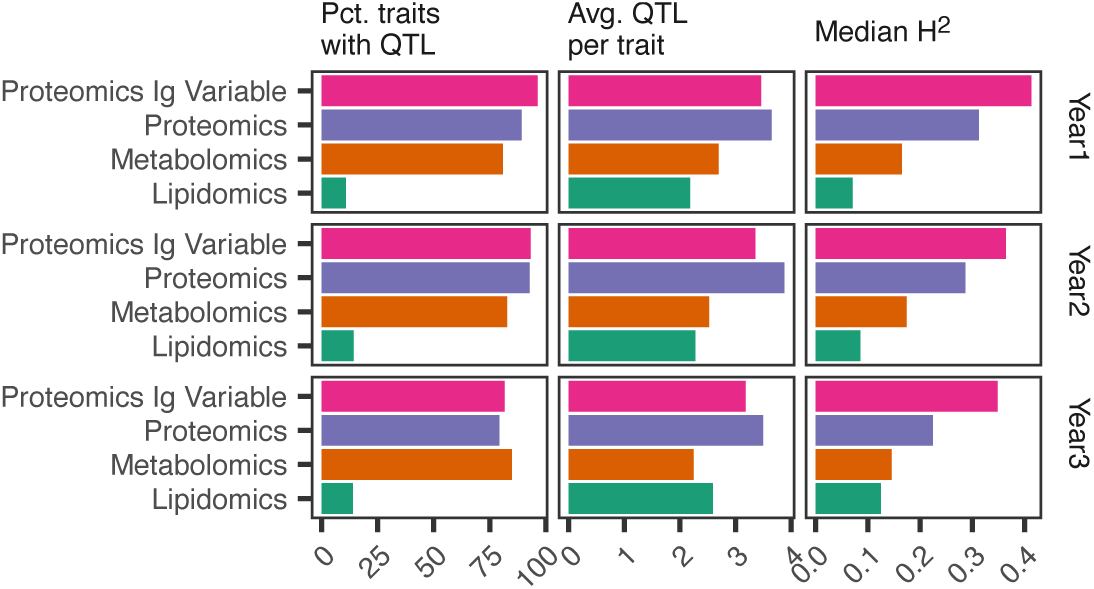
Genetic summary, accounting for immunoglobulin variable regions separately. Percent of traits with QTL, average QTL per trait, and median heritability were calculated separately for compounds annotated as immunoglobulin (Ig) variable regions (Supp. Table 1) versus the rest of the proteins. This was performed due to many genomic coding sites for Ig regions on chromosomes 6 and 12 leading to high number of detected QTLs. Percent of traits with QTL and the average QTL per trait did not differ significantly between Ig variable regions and other proteins.

**Extended Fig. 7.**
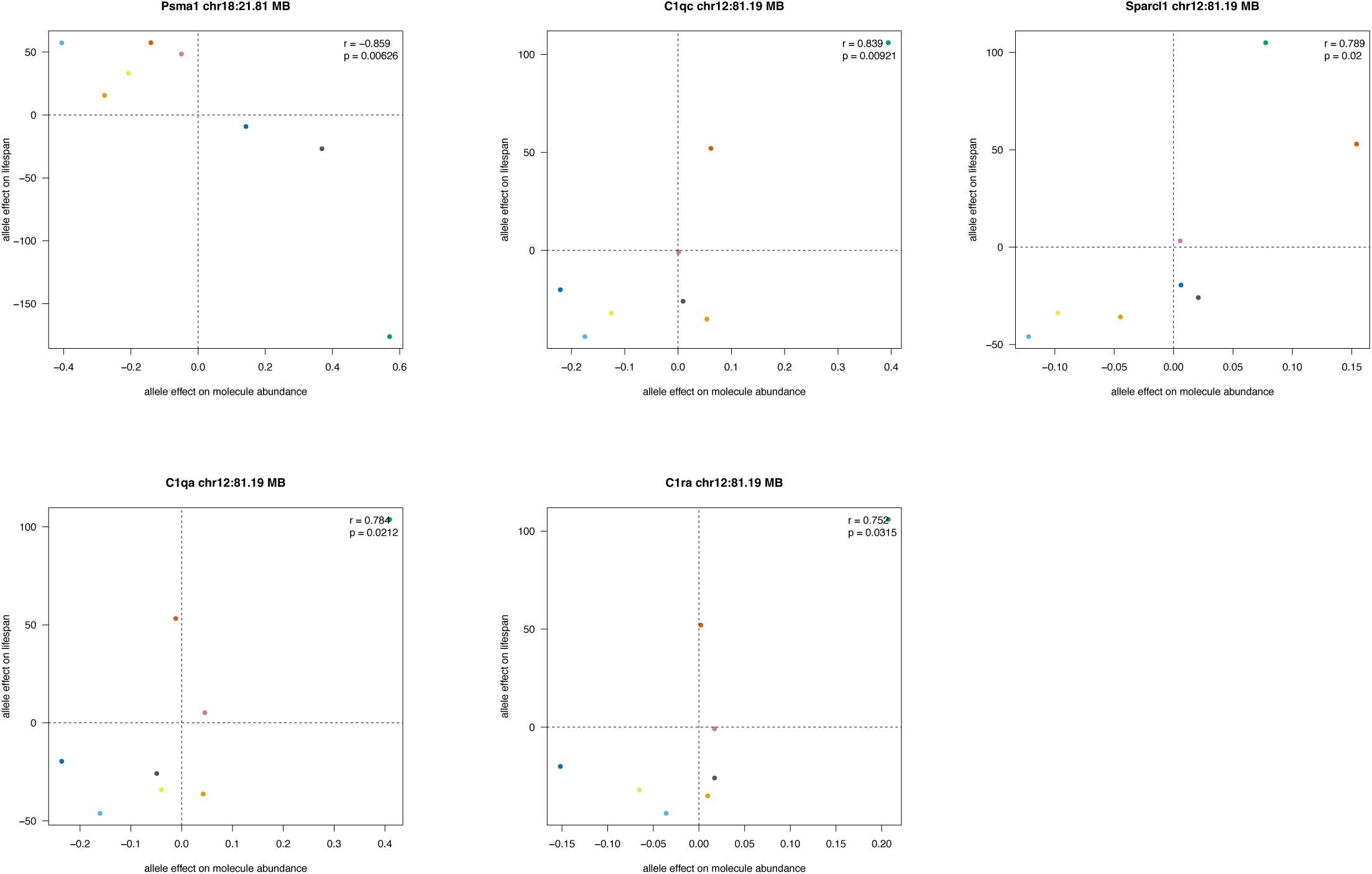
Parent allele affect correlation between molecular abundance and lifespan. Parent allele effect on lifespan and on molecular abundances at identified QTL which colocalized within a 1 MB window of a pre-defined lifespan locus. Correlation performed at chromosomal location of molecular QTL. QTL selected from Figure 5D.

